# Unequal contribution of two paralogous centromeric histones to function the cowpea centromere

**DOI:** 10.1101/2020.01.07.897074

**Authors:** Takayoshi Ishii, Martina Juranić, Shamoni Maheshwari, Fernanda de Oliveira Bustamante, Maximilian Moritz Vogt, Rigel Salinas-Gamboa, Steven Dreissig, Nial Gursanscky, Tracy How, Joerg Fuchs, Veit Schubert, Andrew Spriggs, Jean-Philippe Vielle-Calzada, Luca Comai, Anna M. G. Koltunow, Andreas Houben

## Abstract

The legume cowpea (*Vigna unguiculata*, 2n=2x=22) has significant tolerance to drought and heat stress. Here we analysed and manipulated cowpea centromere-specific histone H3 (CENH3) genes, aiming to establish a centromere-based doubled-haploid method for use in genetic improvement of this dryland crop in future. Cowpea encodes two functional CENH3 variants (CENH3.1 and CENH3.2) and two CENH3 pseudogenes. Phylogenetic analysis suggests that gene duplication of CENH3 occurred independently during the speciation of *V. unguiculata* and the related *V. mungo* without a genome duplication event. Both functional cowpea *CENH3* variants are transcribed, and the corresponding proteins are intermingled in subdomains of different types of centromere sequences in a tissue-specific manner together with the outer kinetochore protein CENPC. CENH3.2 is removed from the generative cell of mature pollen, while CENH3.1 persists. Differences between both CENH3 paralogs are restricted to the N-terminus. The complete CRISPR/Cas9-based inactivation of *CENH3.1* resulted in delayed vegetative growth and sterility, indicating that this variant is needed for plant development and reproduction. By contrast, *CENH3.2* knockout individuals did not show obvious defects during vegetative and reproductive development, suggesting that the gene is an early stage of subfunctionalization or pseudogenization.

**One-sentence summary:** The two paralogous centromeric histones (CENH3) of cowpea contribute unequal to the function of the centromere.

## Introduction

Cowpea (*Vigna unguiculata* (L.) Walp) belongs to the genus *Vigna,* comprising more than 200 species. Cowpea is diploid (2n = 2x = 22) with a genome size of 640.6 Mb (Lonardi et al. 2019). Wild cowpea species are pantropically distributed with highest genetic diversity observed in South Africa, indicating this region is the site of origin (Padulosi and Ng 1997). This herbaceous legume has a pronounced tolerance to drought and heat stress, which allows cultivation on non-irrigated land in semi-arid regions (Hall 2004). Cowpea is one of the eight-grain legumes currently targeted for agronomic improvement by the Consultative Group for International Agricultural Research (CGIAR) (7th CGIAR System Council meeting: https://storage.googleapis.com/cgiarorg/2018/11/SC7-B_Breeding-Initiative-1.pdf). A method to generate doubled-haploids could accelerate the breeding of new, improved, cowpea cultivars. In order to establish a haploidization method based on the manipulation of the centromere (Kalinowska et al. 2019), we analyzed the centromere composition of this species.

The centromeric regions of all cowpea chromosomes are enriched in two repetitive sequences (pVuKB1 and pVuKB2), and seven of the eleven chromosome pairs are additionally marked by a 455 bp tandem repeat (Iwata-Otsubo et al. 2016; Galasso et al. 1999). As centromeric sequences are neither sufficient nor required for centromere identity (Marshall et al. 2008), we focused our analysis on the centromere-specific histone H3 variant CENH3, which is essential for centromere function (Allshire and Karpen 2008). In most diploid eukaryotes and flowering plant species, CENH3 is encoded by a single copy gene. A minority of diploid plants encode two CENH3 homologs including, *Arabidopsis lyrata, Luzula nivea*, *Hordeum vulgare* (barley)*, Secale cereale* (rye), *Pisum sativum* and *Lathyrus sativus* species (Kawabe et al. 2006; Moraes et al. 2011; Sanei et al. 2011; Neumann et al. 2012; Neumann et al. 2015b; Evtushenko et al. 2017). The apparent persistance of two CENH3 paralogs in these species raises the possibility of subfunctionalization, where each has a distinct functional role and which can be tested by studying the effect of individual gene knockouts. A TILLING mutant of the βCENH3 paralog in barley has no phenotype (Karimi-Ashtiyani et al. 2015). However, the barley αCENH3 paralog has not been mutated, therefore functionality could not be evaluated. In tetraploid wheat, virus induced gene silencing (RNAi) used to target both CENH3 types suggested that both paralogs have a functional role, however, RNAi can result in off-target and incomplete silencing effects (Yuan et al. 2015). Therefore, the functional investigation of duplicated CENH3 loci is best evaluated by examining the phenotype of complete CENH3 knockouts.

In this study, we identified two cowpea CENH3 variants, characterized their interaction with the protein CENPC and identified novel centromeric sequences for cowpea. Phylogenetic analyses suggested that the duplication of CENH3 occurred during or before the speciation of *V. unguiculata*. CRISPR/Cas9-based inactivation of both CENH3 variants revealed that CENH3.1 function is required for plant development and reproduction. By contrast, *CENH3.2* knockout individuals did not show obvious defects during vegetative and reproductive development, suggesting that this variant is an early stage of subfunctionalization or pseudogenization.

## Results

### Cowpea encodes two recently evolved functional variants of *CENH3*

*In silico* analysis of the *V. unguiculata* genomic sequence (Phytozome; https://phytozome.jgi.doe.gov/pz/portal.html) resulted in the identification of two *CENH3* variants, which we named: *VuCENH3.1* (Transcript ID: Vigun01g066400) and *VuCENH3.2* (Transcript ID: Vigun05g172200) located on chromosomes 1 and 5, respectively. The intron-exon structure of both *CENH3 genes* is similar, except that the first and second exons of *CENH3.2* are fused (Supplemental figure 1a). The similarity is 91% at the protein level with amino acid differences primarily evident in the N-terminal protein domain (Supplemental figure 1b). Two pseudo genes called *CENH3.3-pseudo* and *CENH3.4-pseudo* (Transcript ID: Vigun01g066300) were also identified incomplete coding regions containing exons 2-4 and 5-7 of *CENH3.1*, respectively (Supplemental figure 1a). *CENH3.3*-*pseudo* is located on chromosome 1 in the promoter region of an unidentified gene (Transcript ID: Vigun01g066200). *VuCENH3.4-pseudo* also encoded by chromosome 1 forms incomplete *CENH3* transcripts (Transcript ID: Vigun01g066300) based on RNAseq analysis (Gursanscky et al. 2019).

To understand the evolution of *CENH3* in cowpea, we analyzed the *CENH3* locus in the draft genomes of legume species *Cajanus cajan*, *Glycine max, Phaseolus vulgaris, V. angularis* and *V. radiata* (Supplemental figure 2a). These analyses indicated that in *G. max* the duplication of *CENH3* arose by whole genome duplication (Neumann et al. 2015a), however, in cowpea the increase in *CENH3* copy number appears to have occurred by duplication at the original *CENH3* locus independent of a whole genome duplication event. *CENH3.3 and 3.4-pseudo* appear to have arisen by tandem gene duplication and pseudogenization (Supplemental figure 2b).

In order to examine if multiple *CENH3* variants exist in other accessions of cowpea and related species, 14 *V. unguiculata* accessions of different origin and nine related cowpea species were examined by analyzing the sequence of RT-PCR products produced using generic CENH3 primers for *Vigna* species (Vigna_CENH3F and Vigna_CENH3R, Supplemental table 1, 2). Two variants of *CENH3* were identified in all *V. unguiculata* accessions, the diploid *V. mungo* and the tetraploid *V. reflex-pilosa*. The diploid *Vigna* species *V. angularis*, *V. umbellate*, *V. aconitifolia*, *V. radiata*, and *V. trilobata*, and the closely related species *V. vexillata* (Takahashi et al. 2016) encode a single *CENH3*. BLAST analysis of publicly available genomic sequence for *V. radiata* (http://plantgenomics.snu.ac.kr/sequenceserver) and *V. angularis* (http://viggs.dna.affrc.go.jp/blast), confirmed that both species encode a single variant of *CENH3*. Alignment of the identified CENH3 amino acid sequences identified in the 23 surveyed *Vigna* species revealed differences in length in the N-terminal domain, however, the length of the histone fold domain remained conserved (Supplemental Figure 3). CENH3 amino acid mutations in *V. unguiculata* accessions containing both CENH3 variants were also found in four positions of CENH3.1 (two in the N-terminal tail, two in the histone fold domain) and three positions of CENH3.2 (one in the N-terminal tail, two in the histone fold domain) (Supplemental Figure 3). Our phylogenetic analysis of *Vigna* CENH3s suggests that duplication of CENH3 occurred independently during or before the speciation of *V. unguiculata* and *V. mungo* (Figure 1).

**Fig. 1.**
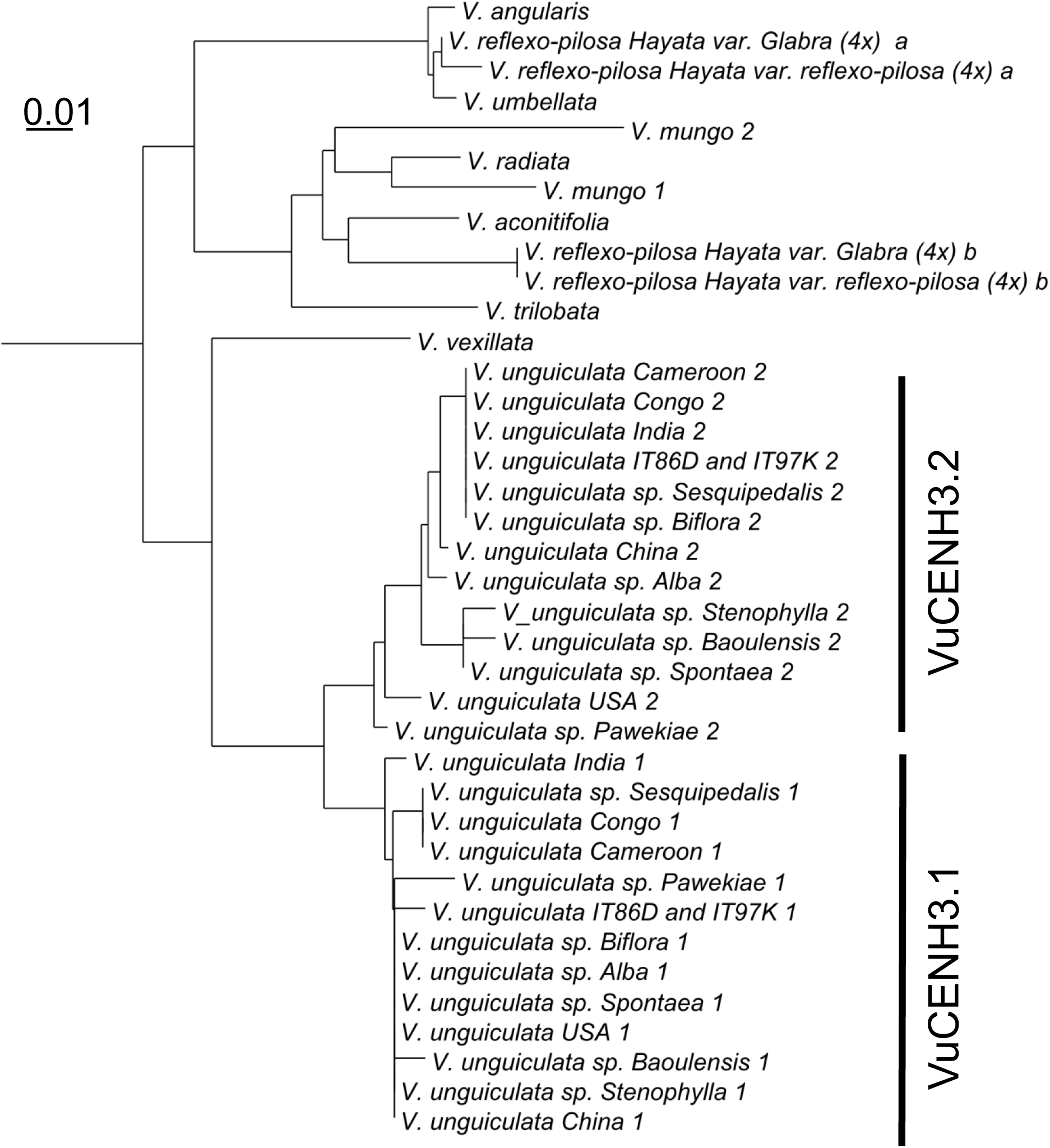
Phylogenetic tree of *Vigna* based on CENH3 amino acid sequences. Two variants of CENH3 (VuCNEH3.1 and VuCENH.2) were identified in all *V. unguiculata* accessions, diploid *V. mungo* and tetraploid species of *V. reflexo-pilosa*. Other diploid *Vigna* species (*V. angularis*, *V. umbellata*, *V. aconitifolia*, *V. radiata*, and *V. trilobata and V. vexillata*) encode a single CENH3.

### *CENH3* variants are transcribed in a tissue-specific manner

The relative expression levels of both functional *CENH3* variants were examined in different cowpea tissue types using quantitative real-time PCR. *CENH3.1* transcripts are more abundant than *CENH3.2* in all tissues analyzed including early and mature anthers, developing carpels, embryos and endosperm of seeds at globular, heart and at cotyledon stages of embryogenesis, leaves, mature ovules, roots and root tips (Supplemental Figure 4a). The highest expression of *CENH3.1* was found in carpel and mature ovule tissue. In addition, RNA-sequencing of laser-captured microdissected (LCM) cells allowed us to understand the *CENH3.1* and *CENH3.2* gene expression in reproductive cell types: the megaspore mother cell (MMC), the tetrad of haploid megaspores, 2- and 4-nuclear embryo sacs, the central cell, the egg cell, as well as the early and late microspore mother cell, the tetrad of haploid microspores, the individual microspore, and the sperm cell (Supplemental Figure 4b). With the exception of the microspore mother cell that showed abundant *CENH3.2* expression at early stages of differentiation, the expression of *CENH3.1* was higher in all other reproductive cells and stages (Supplemental Figure 4b). Transcripts of CENH3.1 were particularly abundant in the MMC, the 2-nuclear embryo sac and the egg cell.

### CENH3.1 and CENH3.2 are co-located in cowpea centromeres

To determine the subcentromeric arrangement of both CENH3 variants in cowpea we generated antibodies against VuCENH3.1 and VuCENH3.2. In addition, an antibody recognizing both variants of CENH3 (anti-VuCENH3 common) was produced (Supplemental Figure 1b). Western blot analysis of nuclear proteins using the developed antibodies revealed CENH3.1 and CENH3.2 dimeric signals, suggesting homodimerization (Supplemental Figure 1c).

Chromosomes were examined by immunostaining to identify locations of CENH3.1 and CENH3.2 proteins. Structured Illumination Microscopy (SIM) analysis showed intermingling of CENH3.1 and CENH3.2 signals in centromeric subdomains of interphase nuclei (Figure 2a), in mitotic chromosomes of root cells (Figure 2b) and also in extended chromatin fibers (Figure 2c). Analyses of multiple extended chromatin fibers isolated from root nuclei indicated that cowpea centromeres comprise intermingled nucleosome clusters containing either one of the two CENH3 variants or CENH3 free nucleosomes. This is consistent with the homodimers detected by Western blotting.

**Fig. 2.**
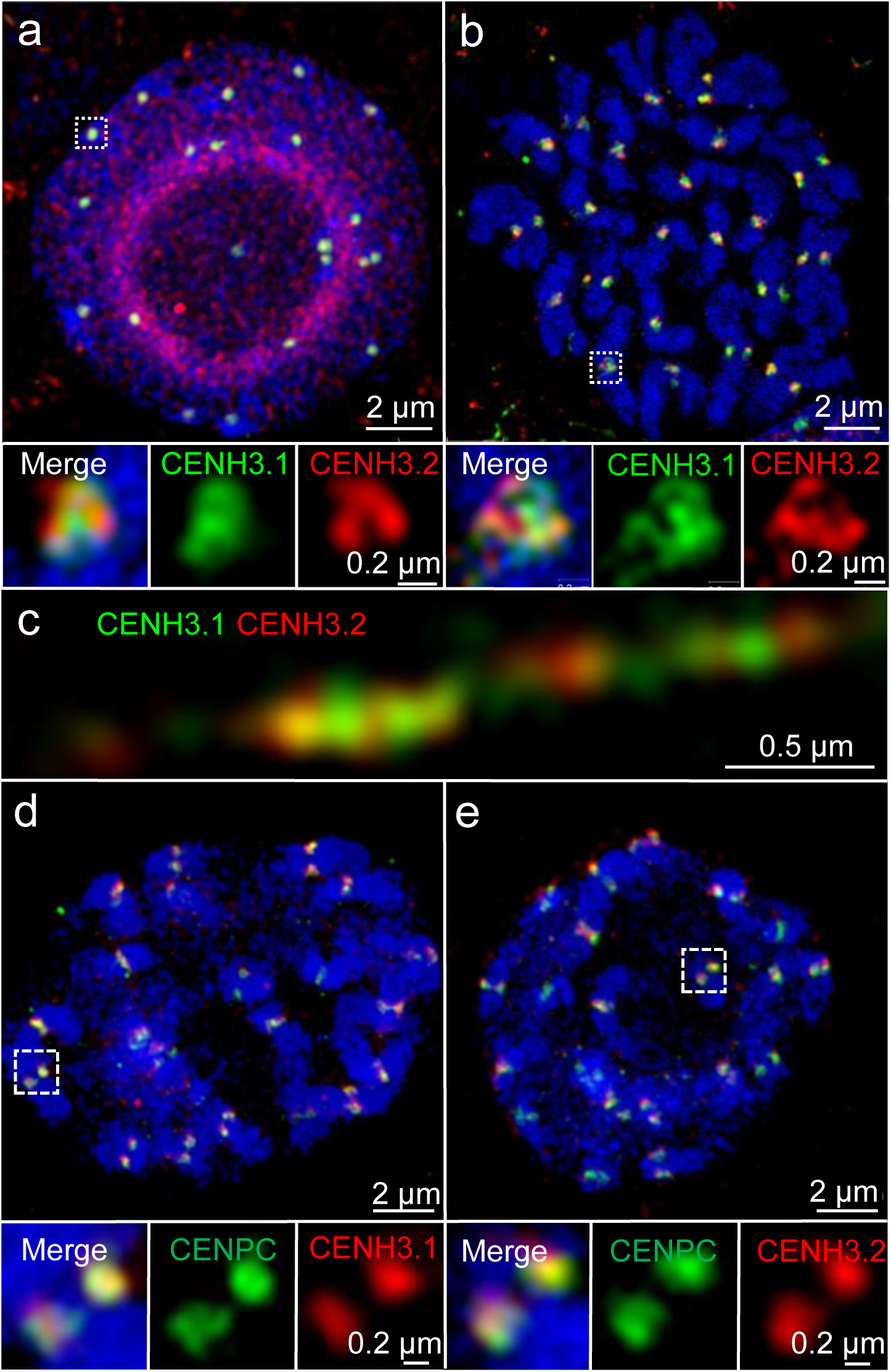
The organization of cowpea centromere analyzed by indirect immunostaining and Structured Illumination Microscopy (SIM) in root cells. Both CENH3.1 (green) and CENH3.2 (red) occupy distinct but intermingled nucleosome domains at centromeres in interphase nuclei (a), prometaphase chromosomes (b), and extended chromatin fibres (c). CENH3.1 (red) and CENH3.2 (red) colocalize with CENPC (green) at the centromeres of prometaphase chromosomes (d and e). Further enlarged centromere regions shown below are indicated (a, b, d and e).

Antibodies to detect cowpea CENPC were generated to provide an additional marker for active centromeres. CENPC is a conserved component of most eukaryotic centromeres that links the inner and outer (microtubule-binding) components of the kinetochore (Earnshaw 2015). CENPC co-localizes with CENH3, defining active centromere chromatin (Marques et al. 2016; Falk et al. 2015; Kato et al. 2013; Carroll et al. 2010). A single *CENPC* candidate (*Vu*CENPC, Transcript ID: Vigun05g287700) was identified in the cowpea genome which aligned with CENPC sequences found in other species (Supplemental Figure 5a). VuCENPC grouped in a sister branch of CENPC sequences identified in other *Vigna* species in phylogenetic analyses (Supplemental Figure 5b). Immunolocalization showed that the CENPC colocalized with immunosignals specific for either CENH3 variant in chromosomes of cowpea roots (Figure 2d,e). In summary, both VuCENH3.1 and VuCENH3.2 protein variants clearly show association with centromeres verifying they are likely to play functional roles in chromosome segregation.

### CENH3 localization dynamics is tissue type-dependent

Next, the distribution of CENH3.1 and CENH3.2 immunosignals was analyzed in nuclei of sporophytic and reproductive tissues to determine the localization patterns of cowpea CENH3s in different tissues in the cowpea plant life cycle. In sporophytic, leaf and root nuclei two different localization patterns of CENH3 were found. A total of 65.9% of leaf nuclei showed centromeric signals for localization of both CENH3s in addition to concomitant nucleoplasmic signals (termed category I). The remaining 34.1% of leaf nuclei (termed category II) showed both CENH3s located only in centromeres (Supplemental Figure 6). By contrast, in roots, 17.6% and 82.2% of nuclei showed category I and II patterns, respectively. Importantly, the similar localization patterns of CENH3.1 and CENH3.2 in these different sporophytic tissue types suggest similar centromere loading of both CENH3 variants.

In contrast to the common behavior in somatic tissues, the two cowpea CENH3s revealed differences when male and female generative tissues were analyzed.

In male meiocytes, both CENH3 variants were found in the centromeres during all stages of meiosis (Supplemental Figure 7). CENH3.1 and CENH3.2 localize to different subdomains of the centromeres at pachytene, metaphase I, and anaphase I chromosomes (Figure 3). By contrast, the loading dynamics of the CENH3 proteins differ during female meiosis. In the female meiocyte (or megaspore mother cell, MMC), whereas CENH3.1 is hardly present during early stages of meiosis I (Supplemental Figure 8), CENH3 is localized in discrete subdomains at leptotene (31.7% of meiocytes), zygotene (32.5%) and pachytene (54.3%) stages, but is absent from adjacent somatic cells in the developing ovule (Supplemental Figure 8). These results indicate that CENH3.2 is the predominantly loaded variant in female meiotic chromosomes.

**Fig. 3.**
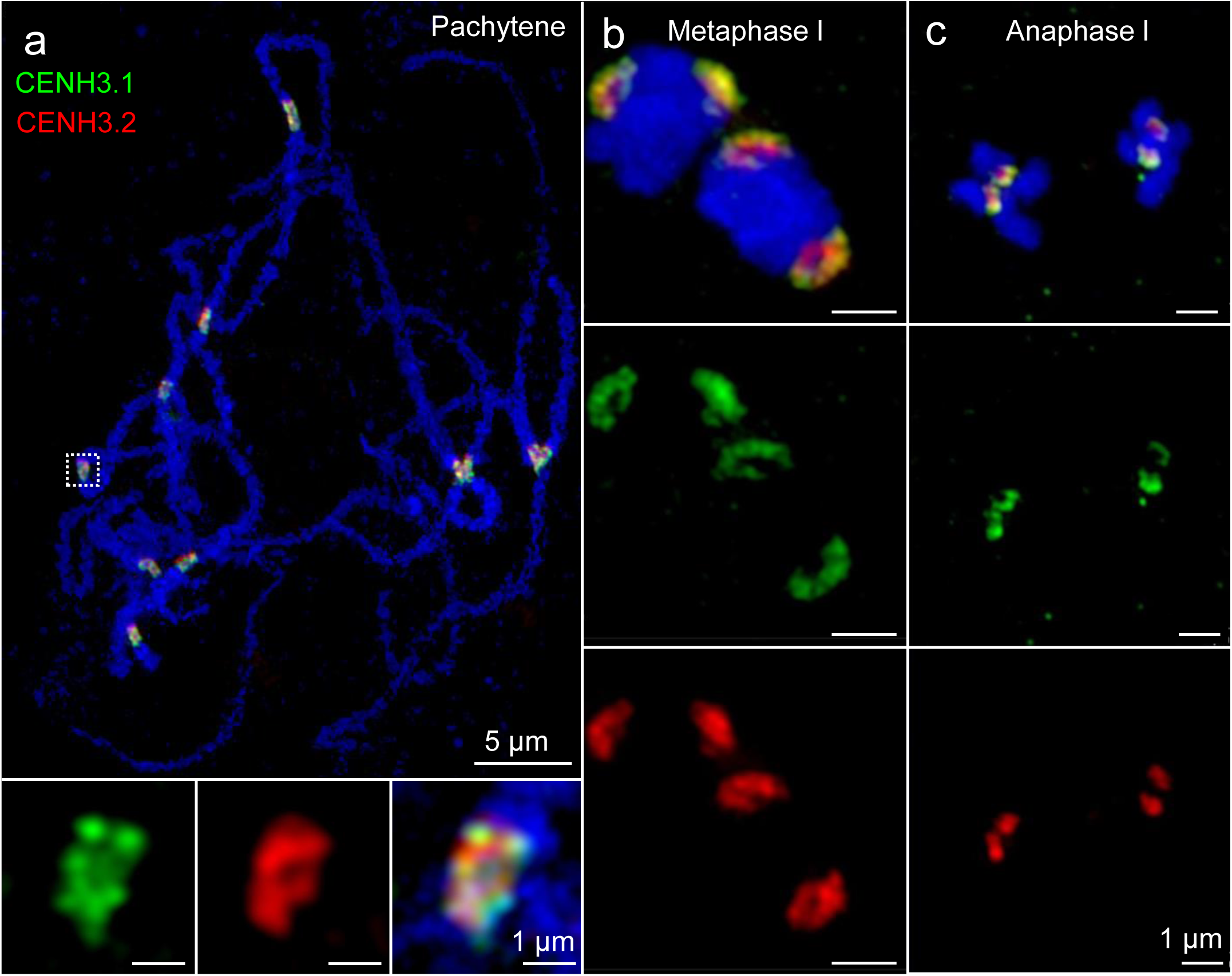
The organization of cowpea centromeres during male meiosis analyzed by indirect immunostaining and Structured Illumination Microscopy (SIM). Both CENH3.1 (green) and CENH3.2 (red) occupy different subdomains of the centromeres at pachytene (a), metaphase l (b) and anaphase l (c) of pollen mother cells. Further enlarged centromere regions of pachytene chromosomes are shown below.

During microgametogenesis, both CENH3 variants marked the centromeres of the unicellular microspore (Figure 4a, b). Notably, in mature pollen, the generative nucleus displayed CENH3.1, but no CENH3.2 signals. As found in *A. thaliana* (Ingouff et al. 2010), the decondensed vegetative nucleus is CENH3-free (Figure 4c, d). The absence of CENPC signals confirms the loss of centromeric proteins in the vegetative nucleus of cowpea (Supplemental Figure 9). Suggesting that CENH3.1 and CENH3.2 are actively removed from the centromeres of the vegetative nucleus. Surprisingly, CENH3.2 is removed from the generative nucleus during the first pollen mitosis. Therefore, in contrast to the similar behavior in vegetative tissue, the two CENH3s display distinct behavior in reproductive tissue.

**Fig. 4.**
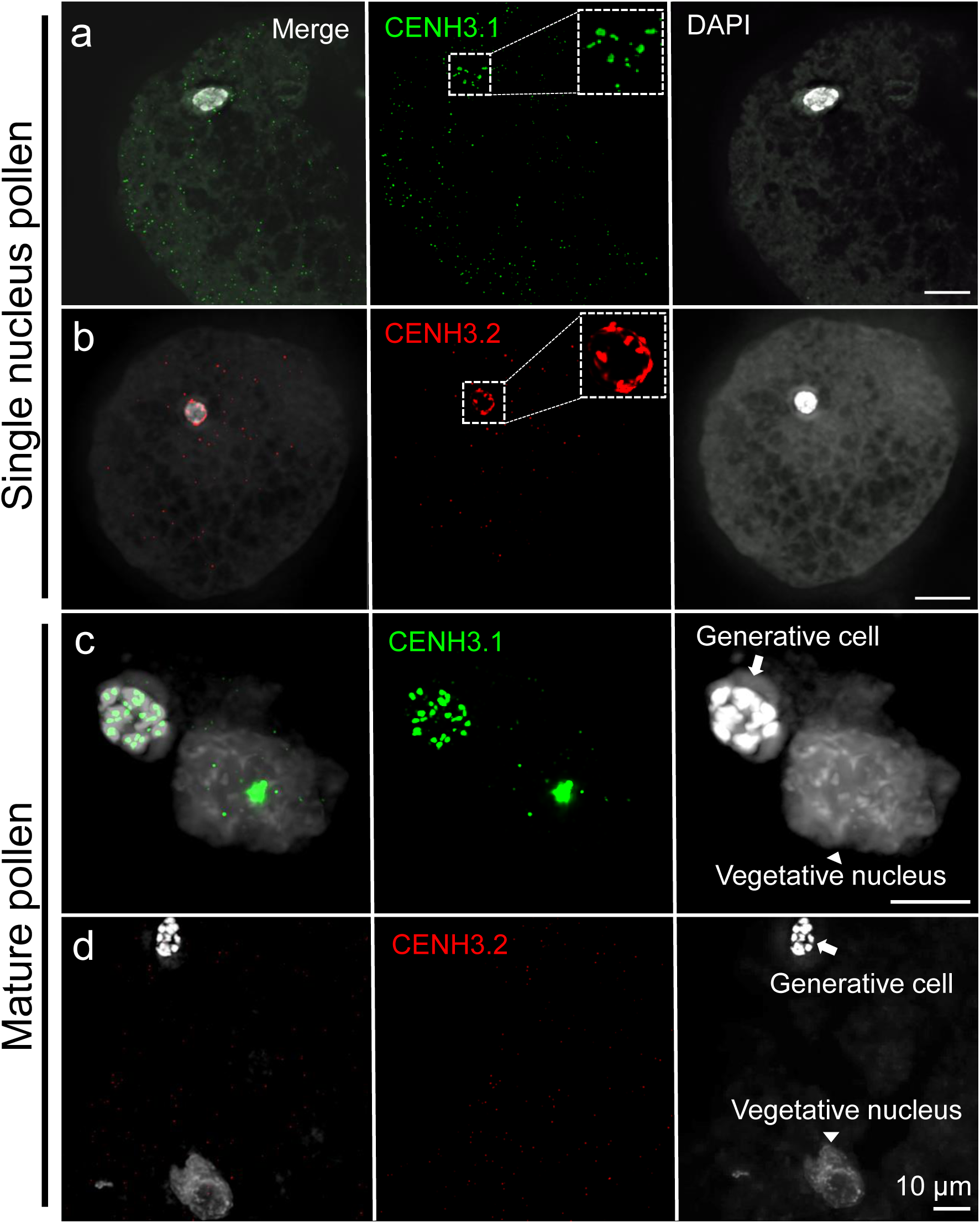
The organization of cowpea centromeres during microgametogenesis. CENH3.1 (green) and CENH3.2 (red) localize in the centromeres of a early-stage mononucleate pollen (a and b). Further enlarged nuclei are shown as inserts. In mature binucleate pollen (c and d), the vegetative nucleus shows no centromeric CENH3.1 (green) and CENH3.2 (red). Centromeric CENH3.1 (green) localizes in the generative nucleus of mature pollen (c), while CENH3.2 (red) does not (d) suggesting specific removal.

Both CENH3 variants co-localized in egg cell centromeres in analyses facilitated using sections of mature ovules from a transgenic cowpea line containing an egg cell specific marker driven by *A. thaliana* DD45 promoter (Table 1, Figure 5). Division of the generative cell into two sperm cells primarily occurs post-pollen tube germination in cowpea (Gursanscky et al. 2019). Following double fertilization both CENH3 variants were observed in centromeres of the immature embryo at the heart stage (Supplement Figure 10). It is possible that after fertilization of the egg cell with the CENH3.2-negative sperm, the centromeres of developing embryos contain both variants of CENH3. Alternatively, *de novo* loading of CENH3.2 occurs at second pollen mitosis post-pollen tube germination. The lack of correlation between transcript abundance and protein localization in both male and female meiocytes suggests that both CENH3 variants are post-transcriptionally regulated in reproductive organs.

**Fig. 5.**
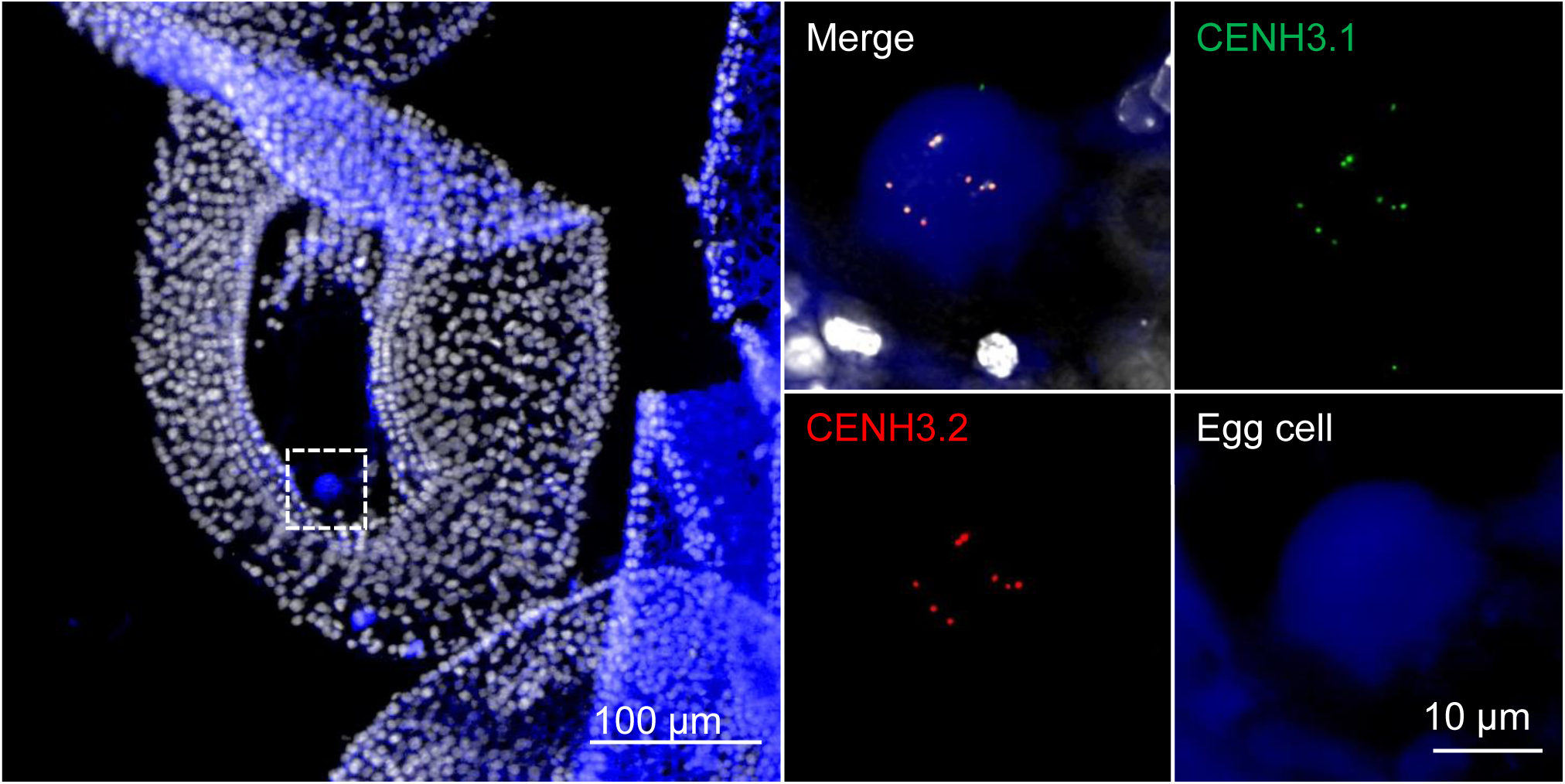
Tissue section of an isolated mature ovule revealing the organization of cowpea centromeres in the egg cell. Both CENH3.1 (green) and CENH3.2 (red) localize at centromeres of the egg cell. The egg cell was identified with an egg cell-specific fluorescence marker (blue), which is driven by the *A. thaliana* DD45 promoter. A further enlarged egg cell region is indicated in the left picture (tissue section of a mature ovule).

### The repeat composition differs between the centromeres of cowpea

Centromeres are often enriched with specific repeats. In agreement with Iwata-Otsubo et al. (2016), pVuKB2-specific signals (Galasso et al. 1999) were found in all centromeres, while only 14 out of 22 centromeres were enriched in 455 bp tandem repeats (Figure 6a). pVuKB2 signals were found to flank the 455 bp tandem repeat in naturally extended pachytene chromosomes (Figure 6b). To determine whether both repeats interact with CENH3-containing nucleosomes and to identify potential additional centromeric repeats in the eight chromosomes found with poor 455 bp repeat labelling, a ChIP-seq analysis was conducted. Two novel centromeric tandem repeats with a repeat unit length of 721 bp and 1600 bp, respectively, were found to interact with CENH3 containing nucleosomes. In addition, the 455 bp tandem repeat (Iwata-Otsubo et al. 2016) also interacted with the CENH3 in nucleosomes thus forming part of the functional centromere. By contrast the pVuKB2 sequence (Galasso et al. 1999) did not associated with CENH3-containing nucleosomes, in line with our FISH data. Both newly identified repeats mark the eight chromosomes found with poor 455 bp repeat labelling (Figure 7a). All three centromeric repeats with a unit length of 455, 721 and 1600 bp, are composed of two to five related sub-repeats, which were named A to E. Unit A is part of all three centromeric repeats (Figure 7b), and shows similarity to Ty3/gypsy retrotransposons, which are often found in plant centromeres (Neumann et al. 2011). No sequence similarity was found between the sequence units A-E and the pericentromeric repeat PvuKB2 (Figure 7c). In conclusion, three repeats are present in CENH3-bound DNA. The 455 bp tandem repeat is dominant in the centromeres of 7 chromosome pairs. The 721 bp and the 1600 bp tandem repeats are major centromere components of the remaining 4 chromosome pairs.

**Fig. 6.**
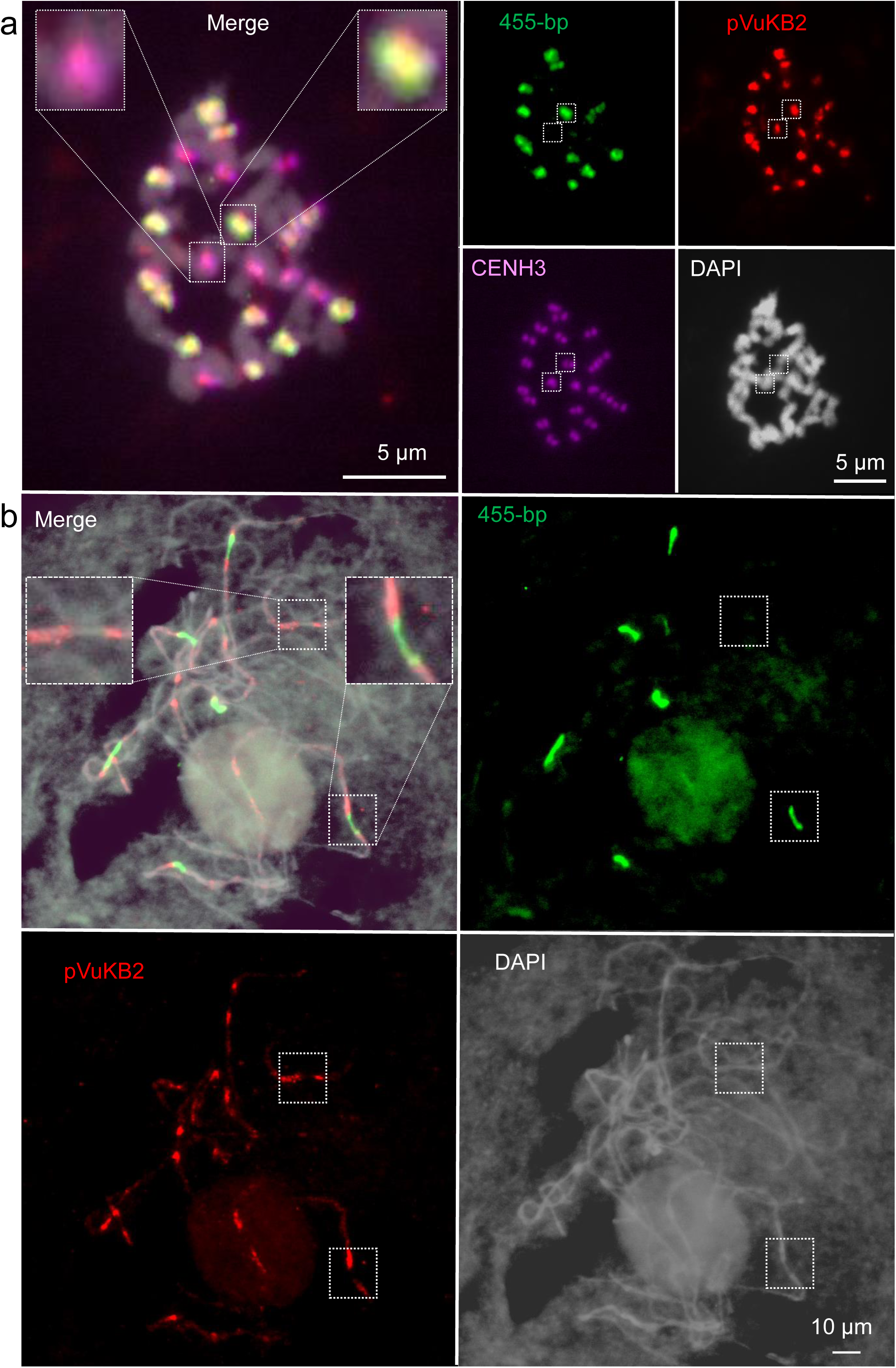
Sequence composition of cowpea centromeres in mitotic (a) and meiotic (b) cells determined by FISH. 14 out of 22 centromeres are enriched in the 455 bp (green) tandem repeat and all centromeres contain the pVuKB2 (red) tandem repeat (a). The position of functional centromeres was confirmed by cowpea CENH3 immunostaining (magenta) (a). pVuKB2 signals (red) are flanked by 455 bp tandem repeats (green) in naturally extended pachytene chromosomes (b). Further enlarged centromere regions are shown as inserts in merged pictures.

**Fig. 7.**
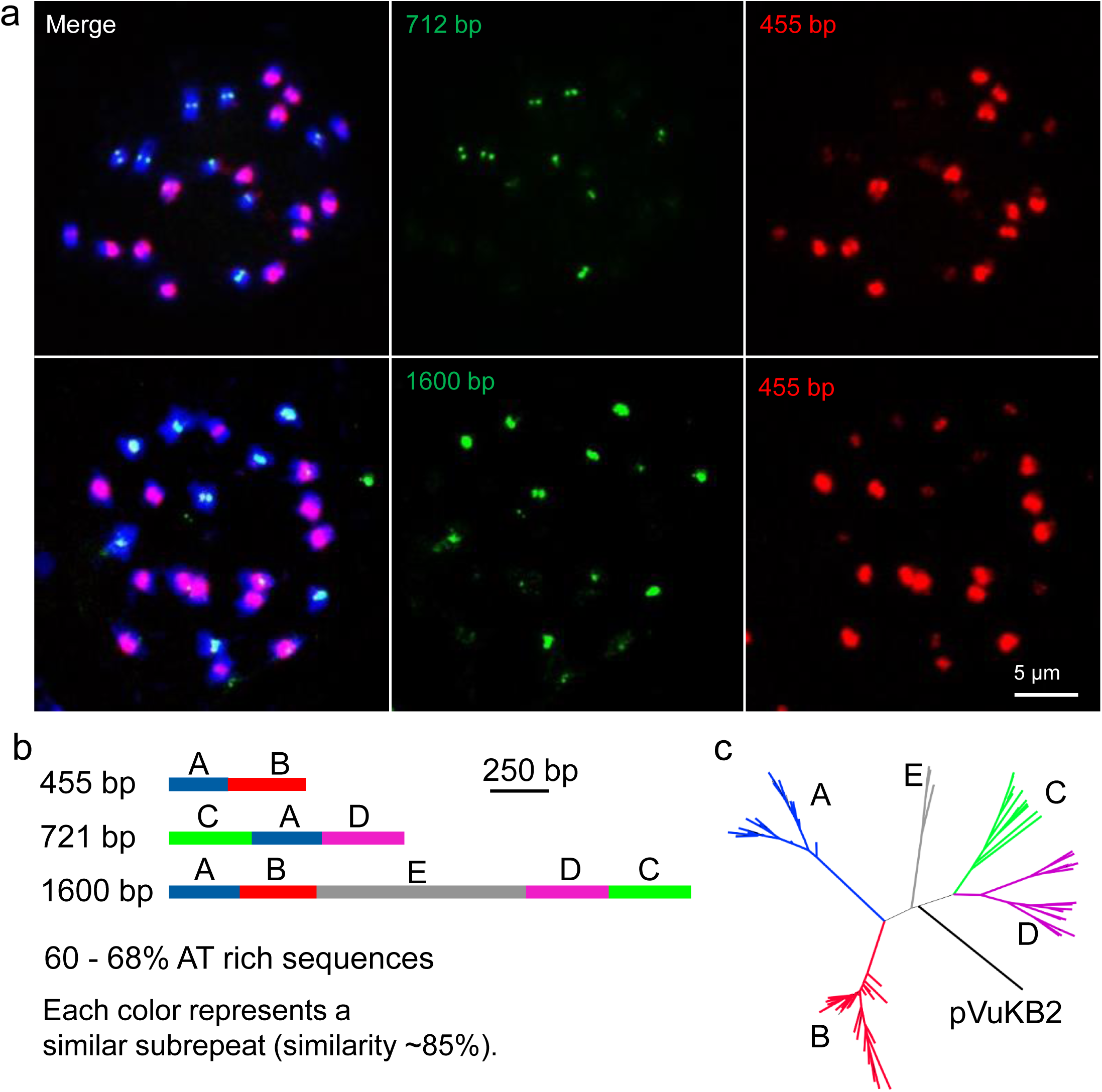
Characterization of novel centromeric tandem repeats of cowpea. Mitotic metaphase chromosomes after FISH with 721 bp (green),1600 bp (red) and 455 bp (red) tandem repeat-specific probes (a). Schematic illustration of the repeat unit (units A – E) organization of 455 bp, 721 bp and 1600 bp centromeric tandem repeats of cowpea (b). Phylogenetic tree based on the DNA sequences of the tandem repeat units A - E and pVuKB2 (c).

### CENH3.1 is sufficient for plant development and reproduction while CENH3.2 is unable to compensate the loss of CENH3.1

CRISPR/Cas9-based genome editing was used to test whether both CENH3 variants are functionally required during cowpea development. Three different guide RNAs were designed to induce mutations in the CENH3 variants. One to induce mutations specifically in CENH3.1 (termed Sg3) and two to induce mutations in both, CENH3.1 and CENH3.2 (Sg4 and Sg5). We generated 19 independent transgenic lines and all were analyzed by TaqMan genotyping. In addition, next-generation (NGS), Sanger sequencing or immunostaining were employed for the characterization of the mutants.

Among 19 T0 plants, four lines had chimeric mutations in *CENH3.1* and two out of these had additional chimeric edits in *CENH3.2*. We focused our analysis on the T0 line named #5B1 (transformed with Sg5), which was mutated in *CENH3.1* (8.7% of NGS reads contained mutations) and *CENH3.2* (37.1% of NGS reads contained mutations) (Supplementary table 3). Further analysis was conducted on T1 progeny of event #5B1. Two of 13 T1 plants (events #5B1-12 and #5B1-13) with chimeric mutations in *CENH3.1* and biallelic mutations in *CENH3.2* were found and both plants were fully fertile (Supplementary table 3). We analyzed ten T2 plants from each event #5B1-12 and #5B1-13, respectively and confirmed the homozygous knockout of CENH3.1 in five T2 plants. A homozygous 1-bp deletion in exon 4 led to a translational frameshift in the CENH3 alpha-N-helix. Among these 20 T2 plants, none had homozygous edits in CENH3.2, but two plants #5B1-12.3 and #5B1-12.4 possessed biallelic heterozygous *CENH3.2* mutations. Finally, we screened for homozygous *CENH3.2* mutations in T3 generation and analyzed twenty T3 plants from events #5B1-12.3 and #5B1-12.4 confirming that six plants were *Cenh3.2 KO* mutant plants while maintaining at least one functional *CENH3.1* allele. All six mutants carried a 2-bp deletion in exon 3, which introduced a stop codon 21-bp downstream from PAM site depleting centromere targeting function of CENH3.2. Immunostaining confirmed the absence of CENH3.1 and CENH3.2 in *Cenh3.1 KO* T2 generation and *Cenh3.2 KO* T3 generation, respectively (Figure 8a). The presence of CENPC in either *Cenh3* mutants suggest that both CENH3 variants interact with CENPC.

**Fig. 8.**
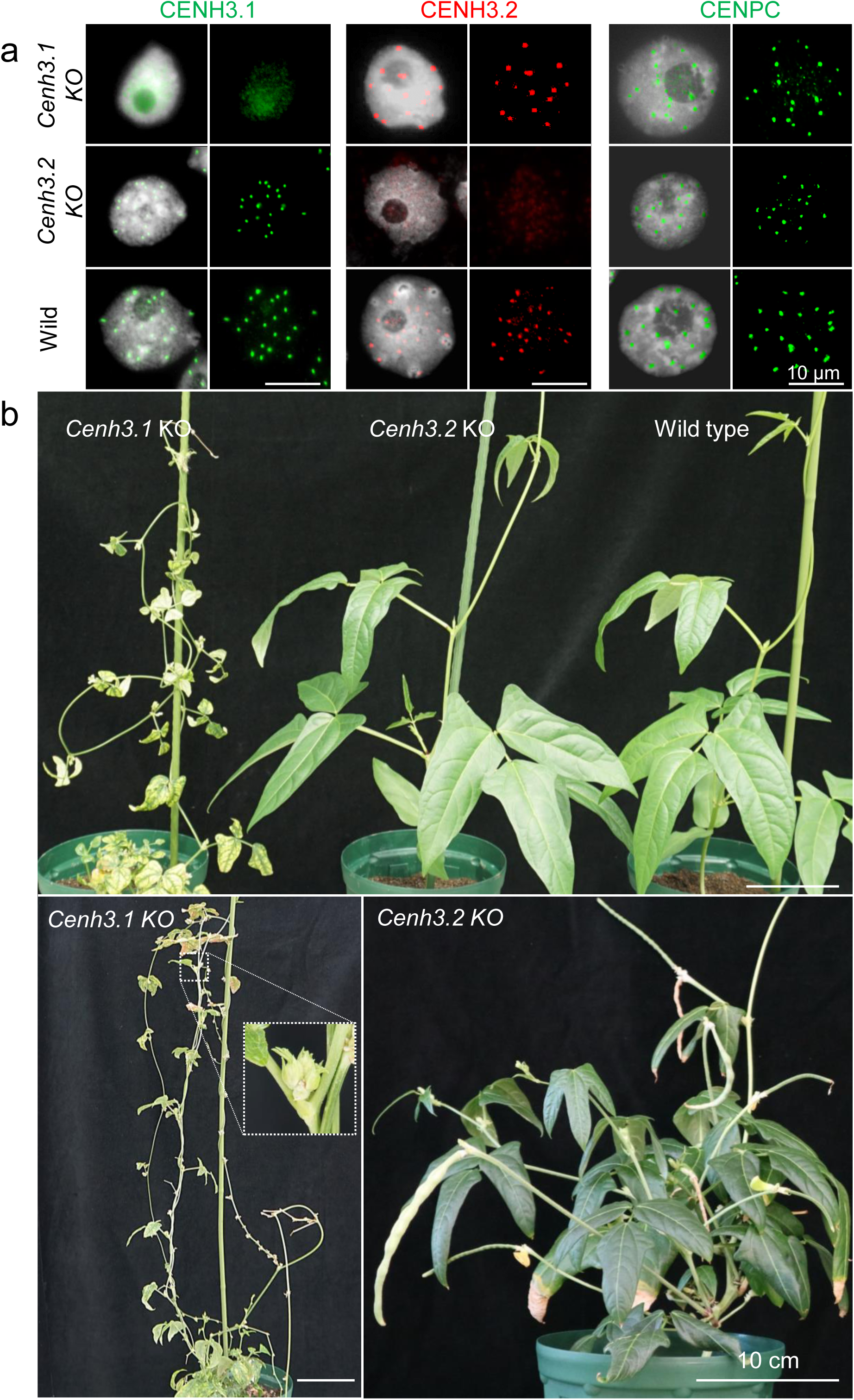
Characterization of *Cenh3.1* and *Cenh3.2* KO plants of cowpea induced by CRISPR/Cas9 based genome editing. Immunostaining of isolated nuclei from *Cenh3.1* and *Cenh3.2* KO plants with anti-CENH3.1 (green), anti-CENH3.2 (red) and anti-CENPC (green) antibodies (a). Plant growth phenotype of *Cenh3.1*, *Cenh3.2* KO and wild-type plants (b). Note the retarded growth of *Cenh3.1* KO plants.

All *Cenh3.1 KO* mutants displayed retarded growth with small necrotic leaves. Flower buds were formed bud stopped development before anthesis (Figure 8b). By contrast, all *Cenh3.2 KO* plants grew similar to the wild type, developed normal flowers and produced normal seed set (Supplementary table 4). Hence, CENH3.1 is essential for normal plant development and CENH3.2 alone, while supporting some growth, is not sufficient for normal development. Moreover, loss of CENH3.2 had no obvious influence on plant growth and reproduction in cowpea under our growth conditions.

## Discussion

### Unusual arrangements of CENH3 genes in *Vigna*

Two CENH3 variants are present in a number of diploid plant species (e.g. *A. lyrata, L. nivea*, *H. vulgare, S. cereale*, *P. sativum* and *L. sativus* species (Kawabe et al. 2006; Moraes et al. 2011; Sanei et al. 2011; Neumann et al. 2012; Neumann et al. 2015a; Finseth et al. 2015; Evtushenko et al. 2017). In animals, multiple copies of *CENH3* have been identified in e.g. *Caenorhabditis elegans*, *C. remanei*, *Bovidae and Drosophila* (Monen et al. 2005; Li and Huang 2008; Monen et al. 2015; Kursel and Malik 2017). In *Drosophila,* the Cid (CENH3) gene underwent at least four independent gene duplication events during evolution of the genus. It has been suggested that retained duplicated CENH3 genes perform nonredundant centromeric functions (Kursel and Malik 2017).

Our analysis of CENH3 in the genus *Vigna* revealed that members of this clade display two alternative genomic configurations: an ancestral one involving a single gene, and one resulting from gene duplication and transposition. We identified two functional CENH3 genes in two diploid species: cowpea and *V. mungo*. In most legumes, such as *P. vulgaris, C. cajan, V. angularis and V. radiata*, the CENH3 locus is syntenic and single copy. A whole-genome duplication in the history of legume evolution dates to 58 Mya (duplication in Papilionoid) (Cannon et al. 2010). However, the presence of only one copy of *CENH3* in *P. vulgaris, C. cajan, V. angularis* and *V. radiata* indicates that loss of one *CENH3* gene occurred after the Papilionoid genome duplication. The presence in soybean of two CENH3 genes at the conserved ancestral position, implies a second whole-genome duplication in soybean (Shoemaker et al. 1996). African *Vigna* (such as cowpea) and Asian *Vigna* (such as *V. angularis, V. radiata and V. mungo*) diverged into different species 4.7 Mya. *V. mungo, V. angularis* and *V. radiata* differentiated 2.8 Mya (Kang et al. 2014). It is likely that ∼4.7 to 2.8 Mya, corresponding to *V. unguiculata* speciation, the ancestral cowpea CENH3 gene on chromosome 10 transposed and duplicated resulting in two loci, one on chromosome 1 and the other on 5 without whole genome duplication. Gene movements could be the result of double-strand break (DBS) repair through synthesis-dependent strand annealing mainly caused by transposable element activity (Wicker et al. 2010). The class II transposons, which are the major group of classical cut-and-paste transposons, comprise 6.1% of the cowpea genome (Lonardi et al. 2019). Chromosome synteny analysis between cowpea and its close relatives *V. angularis, V. radiata* and *P. vulgaris* revealed that chromosome 1 and 5 display rearrangements specific to the genus *Vigna* (Lonardi et al. 2019). In *V. angularis* and *V. radiata*, the ancestral CENH3 locus is conserved suggesting that CENH3 movement only occurred in some *Vigna* species during the rearrangement of chromosomes likely together with the activation of transposable elements. In summary, in the genus *Vigna*, some species contain a single copy of CENH3 while both cowpea and *V. mungo*, have duplicated and transposed genes. When such a case was discovered in *Drosophila* (Kursel and Malik, 2017) it was considered unusual, however, *Vigna,* and others have evolved two CENH3 genes. This poses the question of how duplicated gene copies evolve, and whether they subfunctionalize and are selected, and how they may eventually decay to a single gene configuration.

### Dynamics of CENH3 protein localization in cowpea

We demonstrated that the transcription and centromere occupancy of both cowpea CENH3 paralogs is dynamic and varies among different tissue types. The two types of cowpea CENH3 form intermingling centromeric subdomains in sporophytic (somatic) cell types and in male gametophyte precursor cells undergoing meiosis. A similar subcentromeric organization was reported for the multiple CENH3 variants of *H. vulgare, P. sativum* and *L. sativus* (Ishii et al. 2015a; Neumann et al. 2016). The centromeres of these species are composed of subdomains of either CENH3 variant-containing nucleosome clusters, which, although closely juxtaposed, do not overlap significantly. Due to the restricted optical resolution, it is unclear whether these regions are composed of hetero-nucleosomes containing both CENH3 variants or represent neighbouring CENH3 variants containing homo-nucleosomes as suggested by our Western analysis. The analysis of extended centromeric chromatin fibres revealed the interruption of CENH3-positive nucleosome clusters by clusters of nucleosomes missing CENH3. Thus, cowpea centromeric chromatin fibers seem to be interspersed by nucleosomes containing other histone H3 variants similarly to human and *Drosophila* (Blower et al. 2002; Sullivan and Karpen 2004), suggesting evolutionarily conservation of this arrangement.

The observed centromere organization and dynamics suggests that a CENH3 variant-specific loading is followed by clustering of these nucleosomes into specific centromeric subdomains. In non-plant species, the centromere-targeting domain (CATD) is required for centromere loading of CENH3/CENPA by Scm3/HJURP chaperons (Bassett et al. 2012; Foltz et al. 2009). The CATD domains of both cowpea paralogs are almost identical suggesting that the N-terminal tails, which differ between both CENH3s, are likely involved in the tissue-specific and CENH3 type-specific loading into centromeres.

Subfunctionalization of CENH3 variants was suggested by the expression of cowpea CENH3s during pollen development. In *Arabidopsis*, CENH3 is removed selectively from the vegetative cells (Karimi-Ashtiyani et al. 2015; Merai et al. 2014; Ingouff et al. 2010). As a result, in mature pollen of *A. thaliana* only the sperm nuclei contain CENH3 (Ingouff et al. 2010; Karimi-Ashtiyani et al. 2015). By contrast, the monocotyledonous pearl millet retains CENH3 in the centromeres of both sperm and vegetative cells (Ishii et al. 2015b). In cowpea, at the end of the first pollen mitosis, both CENH3s and CENPC are actively removed from the vegetative nucleus. Unexpectedly, the cowpea CENH3.2 was selectively removed in the generative cell while CENH3.1 was retained and was present in pollen sperms. The differential behavior indicates that a selective removal mechanism recognizes CENH3.2, but not CENH3.1. Given the nearly perfect identity of the histone fold domain of the two paralogs, this implicates the N-terminus in sub functionalization. In contrast to the behavior in pollen, the egg cell retained both CENH3 paralogs. This is the same as found in oat (Ishii et al. 2015b), but different to that described in *Arabidopsis* (Ingouff et al. 2010).

### Contribution of different CENH3 to development

When both CENH3s were knocked out in cowpea via CRISPR/Cas9, both mutant types containing either functional variant of CENH3 displayed vegetative growth, suggesting that both CENH3 paralogs form functional centromeres in somatic tissue. Also, both types of CENH3s are capable of CENPC interaction. However, *Cenh3.1 KO* plants displayed a retarded and abnormal growth phenotype, small necrotic leaves, and incomplete flowers development that did not form seed. In contrast, *Cenh3.2 KO* plants showed normal growth and fertility that could not be distinguished from the wild type. Hence, CENH3.1 of cowpea is essential for normal plant growth and reproduction, while CENH3.2 is a gene likely to be undergoing early subfunctionalization. Its reduced role is consistent with a trajectory of pseudogenization. We cannot rule out, however, that CENH3.2 expression could be advantageous in growing environments that we did not test or that it may contribute to other properties, such as genome stability, that cannot be readily evaluated by observation of two generations. Another possibility is that the inactivation of CENH3.2 during female meiosis results in subtle abnormalities that do not cause female sterility, further cytological analysis of micro- and megasporogenesis in knockout individuals will be necessary to refine the function of CENH3.2 during cowpea reproductive development.

The results in cowpea are consistent with those of barley and indicate that one of the two CENH3 duplicates is dispensable under experimental growing conditions. After inactivation of barley βCENH3, αCENH3 was sufficient for mitotic and meiotic centromere function and development was normal (Karimi-Ashtiyani et al. 2015). The possibility that both CENH3s of cowpea are subfunctionalized and contribute to development is supported by analysis in wheat αCENH3 and βCENH3 (Yuan et al. 2015). Virus induced gene silencing (RNAi) of αCENH3 resulted in extreme dwarfing and a weakened root system suggesting that αCENH3 is essential for plant development. Reduction of plant height and reproductive fitness caused by downregulation of βCENH3 suggest that βCENH3 perhaps plays a more specialized role during reproductive development. Our results are the first to our knowledge to leverage genome editing to understand roles of duplicate CENH3 genes. Consequently, our knockouts are expected to entail null alleles and thus provide firm evidence on the role of individual paralogs.

### The DNA composition of cowpea centromeres

Centromeres are mostly composed of one type of repeat across all chromosomes of a species, such as the α-satellite in human or the 180 bp repeat pAL1 in *A. thaliana* (Murata et al. 1994; Willard and Waye 1987). However, in diploid *Solanum* species, soybean, common bean and chicken, the centromeres are not equally composed and different centromeric sequences exist (Iwata et al. 2013; Gong et al. 2012; Tek et al. 2010; Shang et al. 2010). A similar situation was found in cowpea by immunoprecipitation of CENH3 nucleosomes. The centromeres of this species are composed of three different repeat types. The previously described 455 bp tandem repeat is the major component of the centromeres of 7 chromosome pairs (Iwata-Otsubo et al. 2016). The 721 bp and 1600 bp tandem repeats, identified in this study, are the major centromeric components of the remaining 4 chromosome pairs. Repeat unit A (215 bp) is part of all three centromeric repeats and was identified as a Ty3/gypsy retrotransposon type sequence. None of the centromere repeats of cowpea were found by BLAST analysis in other *Vigna* species. This suggests that the centromere repeat composition in the genus is changing in short evolutionary periods. It will be interesting to further elucidate the evolution of centromeric sequence diversity among different *Vigna* species.

In conclusion, diploid cowpea encodes two types of CENH3. Both functional *CENH3* variants are transcribed and the corresponding proteins are centromere-incorporated in a tissue-specific manner. CENH3.1 and CENH3.2 proteins form intermingling subdomains in all mitotic and meiotic centromeres examined and both CENH3s interact with the binding protein CENPC. Consistent with subfunctionalization, two proteins can show differential expression and localization in cells during plant development. In the most dramatic instance, CENH3.2 is removed from the generative cell of the pollen, while CENH3.1 persists. In the centromeres of seven chromosome pairs of cowpea, CENH3 interacts with the 455 bp tandem repeat, while in the remaining 4 chromosome pairs the centromeres contain the 721 bp and 1600 bp tandem repeats mainly. The centromeric repeats are composed of two to five different subunits, of which only repeat unit A (215 bp) is part of all three centromeric repeats. This repeat unit could be classified as a Ty3/gypsy retrotransposon. Wild-type CENH3.1 is essential for normal plant growth and reproduction, while CENH3.2 is dispensable, suggesting sub-functionalization or pseudogenization of this paralog.

## Methods

### Plant material and growing conditions

The 24 *Vigna* species used in this study (Supplemental Table 1) were germinated and grown in pots (20 cm diameter, 25 cm height) in a greenhouse (16h/8h day-night cycle at 26°C/18°C day-night temperature). Transgenic lines in the *V. unguiculata* cv. IT86D-1010 genetic background were grown under greenhouse conditions in pots (20 cm diameter, 19 cm height) containing Bio Gro^®^ soil mixture (Van Schaik’s Bio Gro Pty Ltd., South Australia) (12h/12h day-night cycle at 28°C/20°C day-night temperature, 40% relative humidity) and a 12h/12h day-night cycle in the Australian Plant Phenomics Facility (APPF), Adelaide.

### Identification of *CENH3* and *CENPC*

CENH3 (Vigan.09G168600) (Sakai et al. 2015; Sakai et al. 2016) of Azuki bean (*Vigna angularis* (Willd.) Ohwi & Ohashi) was used for the *in silico* identification of cowpea CENH3 in genomic and transcriptomic data of cowpea genotype IT97K-499-35 and IT86D-1010 (Spriggs et al. 2018; Lonardi et al. 2019). Trizol-isolated RNA from young leaves were used to generate cDNA with a cDNA synthesis kit (Thermo Scientific). RT-PCR was performed with *Vigna* CENH3-specific primer pairs (Supplemental Table 2). Sequencing of cloned PCR products and amino acid alignments was conducted as described in (Ishii et al. 2015b). CENH3 sequences were submitted to the DDBJ (ID: LC490903 to LC490940).

### Quantitative expression analysis

Total RNA from different tissues of cowpea (mature anther, meiotic anther, carpel, embryo at torpedo stage, leaf, ovule, root, root tip, immature seed at globular and heart stage and whole immature seed) were extracted and used for cDNA synthesis. The absence of genomic DNA was confirmed by PCR using GAPDH-specific primers (Supplemental Table 2). TaqMan-based qRT-PCR was performed in a reaction volume of 10 µl containing 0.5 µl of cDNA, 5 µl of 2×PrimeTime^®^ Gene Expression Master Mix (Integrated DNA Technologies), 0.33 µl (330 nM) primers, 1.25 µl (125 nM) Prime Time locked nucleic acid (LNA) qPCR probes for *CENH3.1* and *CENH3.2* (Integrated DNA Technologies) for increased probe specificity for each gene and *Ubiquitin28* probe for standardization (Eurofins) (Supplemental Table 2). PCR conditions were; 95 °C for 5 min, followed by 35 cycles at 95 °C for 15 seconds and 30 seconds of 61.5 °C using a QuantStudio™ 6 Flex Real-Time PCR System (Thermo Fisher). Three technical replicates were performed for each cDNA sample. Transcript levels of each gene were normalized to *Ubiquitin28* as described in (Ishii et al. 2015a). The specificity and efficiency of all primers were determined by qRT-PCR using a dilution series of cowpea cDNA or cloned CENH3 sequences. Transcript expression patterns for CENH3 genes were also analyzed using LCM-seq datasets based on laser captured microdissection of cowpea reproductive cells as described in (Gursanscky et al. 2019). Reads were aligned against the *V. unguiculata* IT97K-499-35 genome using Biokanga as described (Spriggs et al. 2018; Lonardi et al. 2019) and uniquely aligned reads were counted for each gene.

### Identification of novel centromere repeats

After quality and adapter trimming reads from the ChIP-seq libraries were aligned to a synthetic dimer of the 455 bp repeat using bwa (Li and Durbin 2009) with default settings to estimate the efficiency of the VuCENH3 ChIP. The dataset with the greatest enrichment of the 455 bp repeat (29.6%) was selected and depleted of reads that aligned to the reference. The remaining reads were clustered using CDHIT-EST (Li and Godzik 2006) at 0.9 sequence identity threshold. The ChIP and input datasets were mapped to representative sequences from these clusters as contigs and DESeq(Anders and Huber 2010) analysis was performed to identify contigs enriched in the VuCENH3 ChIP. The enriched sequences were mapped back to the cowpea genomic contigs (IT86D_1010), of which a majority overlapped tandem repeats, with a periodicity of 721 (DDBJ ID: LC490941) and 1600-bp (DDBJ ID: LC490942).

### Indirect immunostaining and Western analysis

The following peptides were used for the production of polyclonal antibodies in rabbits (VuCENH3.1: PASLKVGKKKVSRASTSTP, VuCENH3.2: ASLKASRASTSVPPSQQSP, VuCENH3 common: QQSPATRSRRRAQEEEPQE and VuCENPC: RPVYGRIHQSLATVIGVKCISPGSDGKPTMKVKSYVSDQHKELFELASSY). LifeTein (www.lifetein.com) and Li International (www.liinternationalbio.com) performed the peptide synthesis, immunization of rabbits, and peptide affinity purification of antisera. CENH3 antibodies were directly labelled with Alexa fluor 488 NHS ester (Thermo Fisher) or NHS-rhodamine (Thermo Fisher) as described (Maheshwari et al. 2017). Mature ovules were fixed in 1× phosphate-buffered saline (PBS) containing 4% paraformaldehyde (PFA) under vacuum at 4 °C for 10 min followed by a 5 h fixation at 4 °C without vacuum. Fixed cowpea ovules were embedded and sectioned according to (Wu et al. 2019). For the analysis of pollen mother cells, immature anthers were fixed with 1× PBS containing 4% PFA under vacuum at 4 °C for 10 min followed by a 30 min fixation at 4 °C without vacuum. The anthers were washed with ice-cold 1× PBS for 3 min two times, and digested with an enzyme cocktail composed of 1% (w/v) pectolyase (Sigma), 0.7% (w/v) cellulase ‘ONOZUKA’ R-10 (Yakult), 0.7% cellulase (CalBioChem), and 1% cytohelicase (Sigma) dissolved in 1x PBS for 30 min at 37 °C in a humid chamber. Anthers were subsequently washed with ice-cold 1× PBS for 3 min two times. Excised pollen mother cells were squashed in 1× PBS between slide and coverslip. Slides were used for immunostaining after removing the coverslips. Chromosome spreads derived from root meristems and mature pollen and chromatin fibres for immunostaining were processed as described in (Maheshwari et al. 2017; Ishii et al. 2015b). Whole-mount protein immunolocalization in female meiosis was performed as previously described (Salinas-Gamboa et al. 2016). Primary antibodies were used at a dilution of 1:100. Secondary antibody was Alexa Fluor 488 (Molecular Probes) at a 1:300 dlution. After secondary antibody incubation, slides were treated with propidium iodide in 1× PBS for 20 min, washed for 30 min in 1× PBS, and mounted in PROLONG medium (Molecular Probes). Serial sections were captured on a confocal laser scanning microscope (Zeiss LSM 510 META), with multitrack configuration for detecting iodide (excitation with a diode-pumped solid-state laser at 568 nm, emission collected using a band-pass of 575 to 615 nm) and Alexa 488 (excitation with an argon laser at 488 nm, emission collected using a band-pass of 500 to 550 nm). Laser intensity and gain were set at similar levels for all experiments. Western blotting analysis was performed as described in (Karimi-Ashtiyani et al. 2015).

### Native chromatin immunoprecipitation (ChIP-seq)

Nuclei were isolated from roots and leaves according to (Gendrel et al. 2005) from 3 - 4 day old cowpea seedlings grown at 26 °C in darkness. Nuclei in MNase buffer were digested with 0.5 gels U/μl micrococcal nuclease (NEB) for 25 min at 37 °C. The reaction was stopped with 50 mM EDTA, and the S1 and S2 chromatin fractions were prepared as described in (Maheshwari et al. 2017). For ChIP experiments, 700 µl of the combined S1 and S2 solution was adjusted to a final volume of 2 ml using the ChIP dilution buffer (39 mM NaCl, 20 mM Tris HCl pH 8.0 and 5 mM EDTA). 10 µg of antibody was bound to Dynabeads Protein A (Invitrogen) following manufacturer guidelines at 4 °C for 4 h. Antibody-coated Dynabeads were mixed with 2 ml ChIP solution and incubated overnight at 4 °C using a rotating shaker. Immuno-precipitated complexes were processed accordingly (Maheshwari et al. 2017). Immunoprecipitated DNA and input samples were used for library preparation following manufacturer’s recommendations (Illumina TruSeq ChIP Sample Preparation Kit #IP-202-1012). Subsequently, prepared libraries were single-end sequenced 100 bp on Illumina HiSeq 2000. The original ChIP-seq sample data are available under study accession number PRJEB9647 at the EBI data base (http://www.ebi.ac.uk/ena/data/view/PRJEB33419).

### Fluorescence *in situ* hybridization (FISH)

Root tips were pretreated with 2 mM 8-hydroxyquinoline at room temperature for 4 h. Then, the material was fixed with 6:3:1 (V/V) ethanol/chloroform/glacial acetic acid for 3 days and stored at 4 °C until use. Slide preparation and FISH was performed as described in (Ishii et al. 2010). Specific sequence in pVuKB2 (Galasso et al. 1999), 721-bp tandem and 1600-bp tandem repeats were selected and PCR labelled with tetramethyl-rhodamin-5-dUTP (Roche) (Ishii et al. 2010). In addition, cyanine 5 5’ labelled 20 nucleotide-long oligos (Operon) were used as FISH probes for 455-bp repeat (Supplemental Table 2).

### Microscopy

Standard fluorescence microscopy and Structured Illumination Microscopy (3D-SIM) analysis were performed as described in (Maheshwari et al. 2017). Whole-mount immunostaining serial sections of stage 1 ovules were captured on a laser scanning confocal microscope (Leica), with multitrack configuration for detecting PI (excitation with laser at 568 nm, emission collected using BP: 575–615 nm) and Alexa 488 (excitation with Argon laser at 488 nm, emission collected using BP: 500–550 nm). Laser intensity and gain were set at similar levels for all experiments, using negative controls to adjust them and avoid overexposure and autofluorescence. Projections of selected optical sections were generated using Photoshop.

### Plant transformation

For CRISPR/Cas9-based gene editing, guide RNAs were designed with CRISPRdirect (Naito et al. 2015), cloned into pChimera and into the binary vector pCAS9-TPC (Fauser et al. 2014). Guide RNAs used Sg3: a CENH3.1-specific SgRNA CTGCGACAAGAAGTCGTAGA-PAM; Sg4 and Sg5: targeting both CENH3s GCTCAAGAAGAGGAGCCGCA-PAM, and GCAGCAGCGCCACAGACTCA-PAM, respectively. Final constructs were electroporated into the *Agrobacterium tumefaciens* strain AGL1 for use in cowpea stable transformation according to (Popelka et al. 2006). Transformants were selected on medium containing 2.5 mg/L Basta (Hoechst) herbicide. Shoots developing healthy roots were transferred into 90 mm small pots containing sterilized soil mixture (Van Schaik’s Bio-Gro Pty Ltd, Australia), acclimatized in the growth room at 22°C with 16 h photoperiod for up to 4 weeks and then transferred to the glasshouse in larger pots. PCR was performed to confirm the presence of the *Cas9*, *pat* and *gRNA* genes with the primers listed in Supplemental Table 2. Generation of a fluorescent transgenic reporter line, which carries an egg cell-specific promoter AtDD45 was described in (Gursanscky et al. 2019).

### Analysis of genomic edits in cowpea transgenic lines

DNA extracted from leaf tissue of transgenic T0 plants carrying CRISPR/Cas9 T-DNA was used for Illumina Amplicon-MiSeq DNA sequencing. Target regions spanning the Cas9/sgRNA target site of *CENH3.1* and *CENH3.2* genes were PCR amplified using primers listed in Supplemental Table 2. Amplicons were submitted for 150 PE sequencing on the Illumina MiSeq platform at the Australian Genome Research Facility (AGRF, Melbourne). Mutations induced at the protospacer sites were analyzed with CRISPR RGEN Tools Cas-Analyzer software (Park et al. 2017). Target regions were also amplified from transgenic T1 and T2 plants and cloned into pCR^®^-Blunt II-TOPO^®^ vector (Invitrogen) for analysis by Sanger sequencing.

### TaqMan-based genotyping

TaqMan-based genotyping of plants was performed as described in (Findlay et al. 2016). Briefly, 5 µl of 2x PrimeTime® Gene Expression Master Mix (Integrated DNA Technologies), 0.33 µl (330 nM) of forward and reverse primers (Supplemental Table 2), 1.25 µl (125 nM, Supplemental Table 2) of TaqMan®-Probes (Drop off probe and Reference probe), 1 ul (50 ng/ul genomic DNA) 1.59 µl of water using the following conditions with: 95 °C for 5 min, followed by 35 cycles at 95 °C for 15 seconds and 30 seconds of 69 °C (Ramp rate with 0.8 °C /s decrease the temperature) and end-read of the fluorescence and plot the fluorescence intensity with scatter chart using a QuantStudio™ 6 Flex Real-Time PCR System (Thermo Fisher).

## Acknowledgements

*Vigna* species seeds were kindly provided by Botanic Garden Meise in Belgium. IITA for providing IT86D-1010 and IT97K-499-35 cowpea lines for use in the research. Thanks to Natalia Bazanova, Dilrukshi Nagahatenna (CSIRO), Jana Lorenz and Sylvia Swetik (IPK) for technical assistance and maintenance of transgenic plants and Jennifer Taylor (CSIRO) for bioinformatics advice and support. Thanks to Axel Himmelbach (IPK) for Next-Generation Sequencing and Anne Fiebig (IPK) for submission of sequence data to the European Nucleotide Archive.

## Funding

This work was supported by a sub-award from the CSIRO for the grant ‘Capturing Heterosis for smallholders: OPP1076280’ from the BMGF (USA).

## Authors contributions

T.I., M.J. and A.H. designed the experiments. T.I., M. J., S.M., F.O.-M., M.V., R.S.-G., S.D., N.G., T.H., J.F., V.S., and A.S. performed the experiments. T.I., M.J., L.C., J. V.-C., A.K. and A.H. wrote the paper.

## Supplementary information

**Fig. S1.**
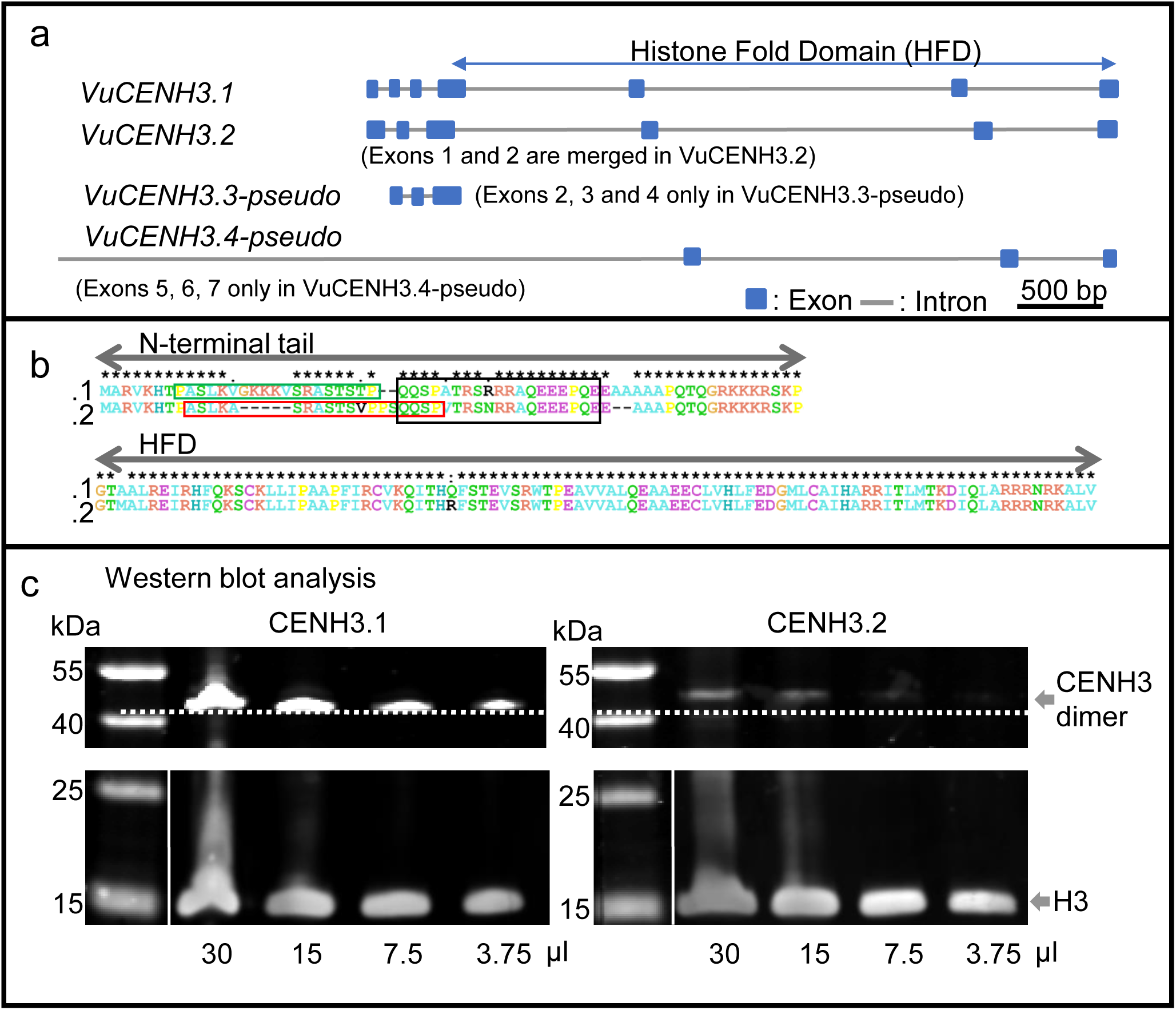
Schematic illustration of the cowpea CENH3 gene exon and intron structure (a). Alignment of CENH3.1 and CENH3.2 amino acid sequences. Green, red and black boxes indicate the position of the peptides used for the generation of CENH3.1, CENH3.2 and CENH3 common antibody, respectively (b). Western blot analysis of cowpea nuclear proteins with CENH3.1-, CENH3.2- and histone H3-specific antibodies. Different loading amounts of nuclear proteins are indicated (30 - 3.75 μl) (d).

**Fig. S2.**
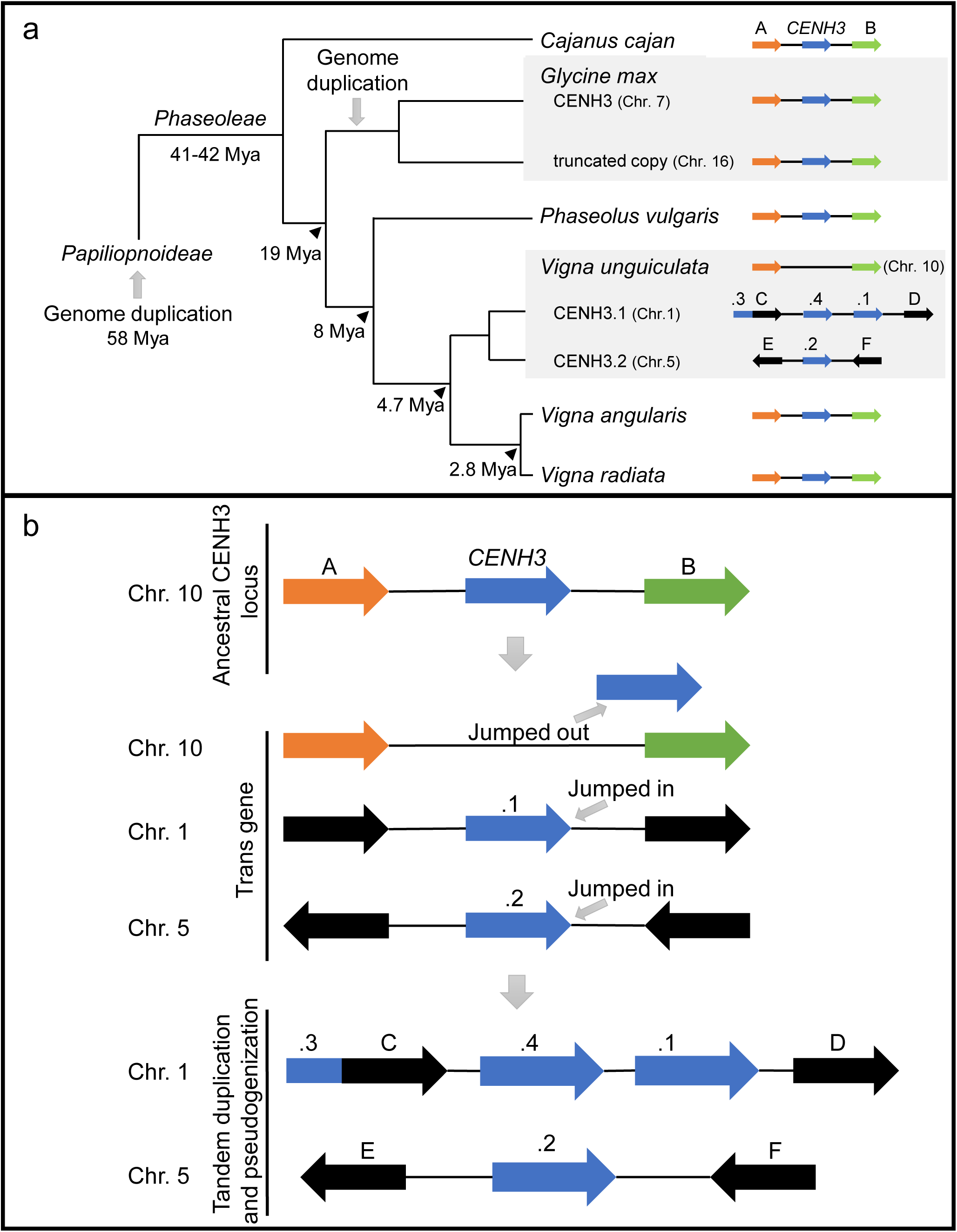
Phylogenetic tree based on the CENH3 amino acid sequence, with a scheme of the chromosomal *CENH3* gene locus on the right of each species. Genome duplication events and node ages are based on (Lavin et al. 2005; Cannon et al. 2010; Kang et al. 2014) (a). Schemata of possible mechanism of CENH3 duplication in cowpea genome (b). Blue arrows indicate *CENH3* gene locus and nearby genes indicated by different colours as A: Rhodanese-like domain-containing protein, B: WRKY transcriptional factor, C: unknown (exosome complex exonuclease), D: unknown, E: calcium-dependent lipid-binding domain-contained protein, F: 60S ribosomal protein L18A, respectively.

**Fig. S3.**
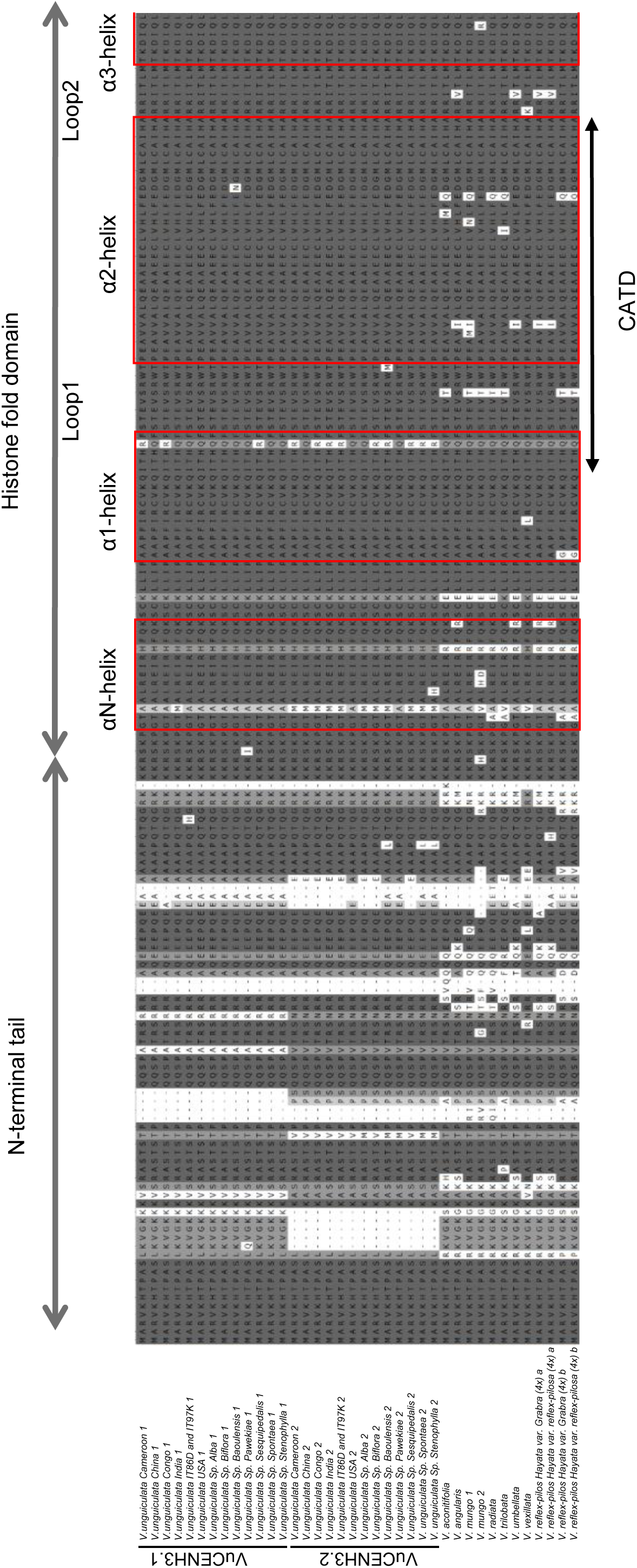
Diversity of CENH3 in *Vigna* species. Multiple alignments of CENH3 proteins from different cowpea accessions (*V. unguiculata -*Cameroon, - China, -Congo, -India, -IT86D-1010, -IT97K-499-35 and -USA), different subspecies of *V. unguiculata* (Alba, Biflora, Baoulensis, Pawekiae, Sesquipedalis, Spontanea, and Stenophylla), different diploid *Vigna* species (*V. aconitifolia, V. angularis, V. mungo, V. radiata, V.trilobata, V. umbellate* and *V. vexillata*), and tetraploid *Vigna* species (*V. reflexo-pilosa var glabra* and *V. reflexo-pilosa var. reflexo-pilosa*). Conserved CENH3 domains are indicated with red-boxes.

**Fig. S4.**
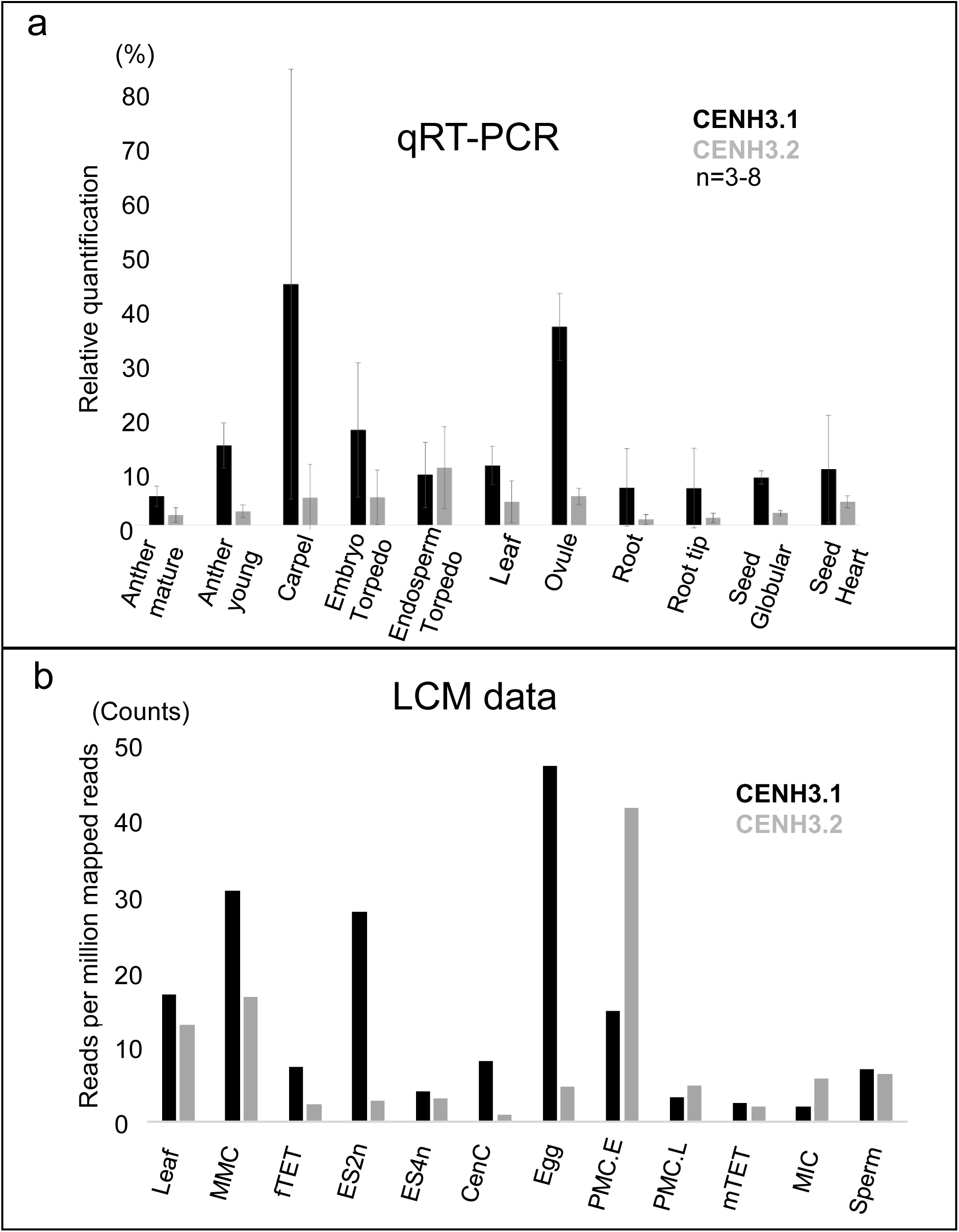
Gene expression patterns of CENH3.1 and CENH3.2 in different tissue and cell types of cowpea. qRT-PCR analysis using RNA isolated from different tissues of cowpea (a). RNA-sequencing using RNA isolated from laser capture microdissected cell types of cowpea (b). Leaf, MMC-megaspore mother cell, fTET-female tetrads, ES2n-embryo sac (2 nuclei), ES4n-embryo sac (4 nuclei), CenC-central cell, egg, PMC.E-early pollen mother cell, PMC.L-late pollen mother cell, mTET-male tetrads, MIC-microspore, sperm.

**Fig. S5.**
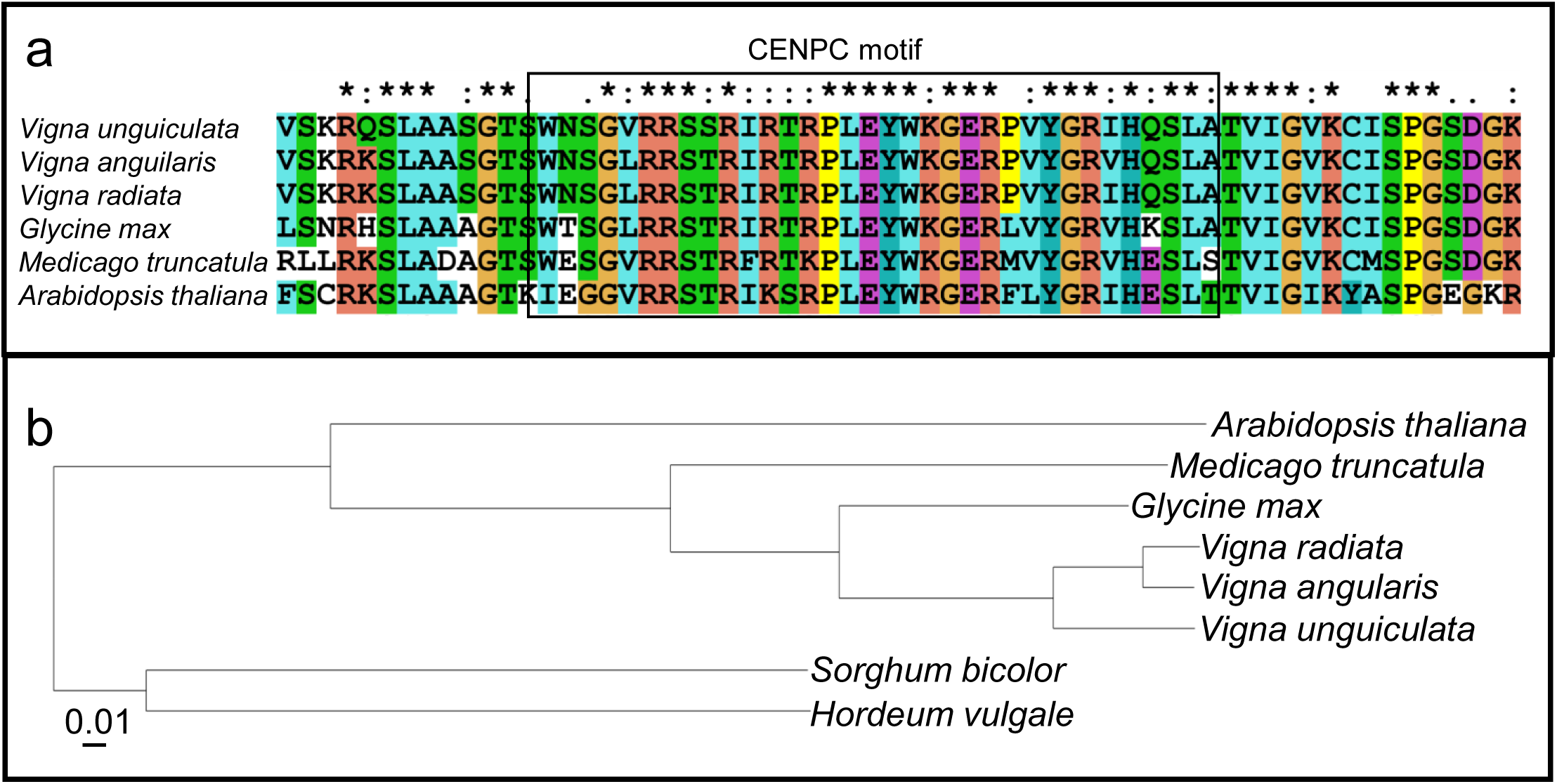
Identification of cowpea CENPC. Alignment of partial CENPC proteins of *A. thaliana, G. max, M. truncatula, V. angularis, V. radiata* and *V. unguiculata* (a). Phylogenetic tree based on the full length CENPC proteins of *A. thaliana, G. max, H. vulgale, M. truncatula, S. bicolor, V. angularis, V. radiata* and *V. unguiculata*.

**Fig. S6.**
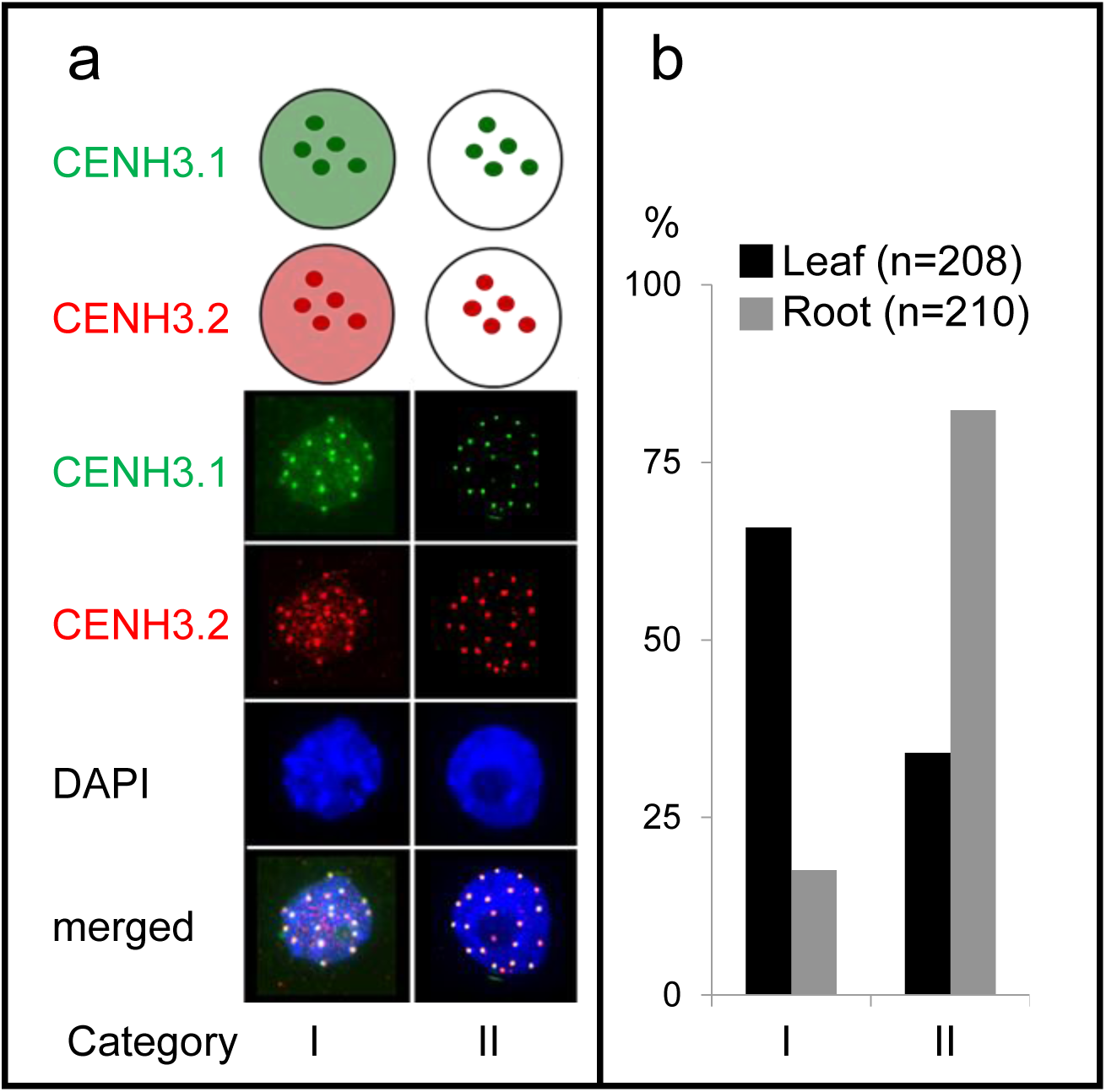
Distribution of CENH3.1 and CENH3.2 immunosignals in isolated nuclei from leaf and root tissues of cowpea. Two categories are determined based on the configurations of CENH3 signals (a). Category l: Centromeric CENH3.1 and CENH3.2 signals colocalize and nuclei reveal disperse immunosignals. Category ll: Centromeric CENH3.1 and CENH3.2 signals colocalize without dispersing signals. Typical examples are shown in (a). Quantification of categorized leaf (n=208) and root (n=210) nuclei (b). Black bar: leaf nuclei, grey bar: root nuclei.

**Fig. S7.**
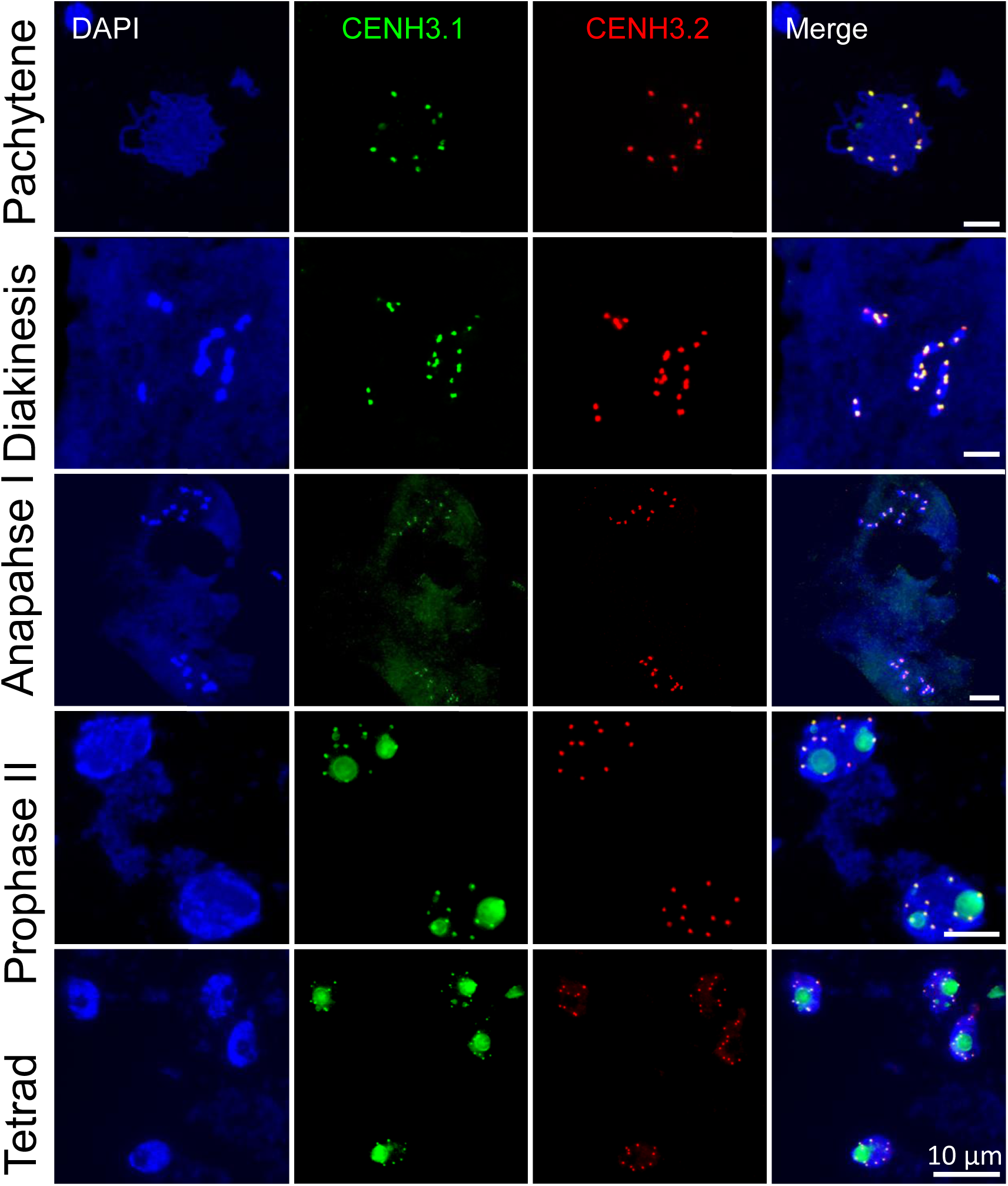
Centromeric CENH3.1 and CENH3.2 colocalize during all stages of male meiosis in cowpea. Distribution of CENH3.1 (green) and CENH3.2 (red) are shown in pachytene, diakinesis, anaphase l, prophase ll and tetrad cells.

**Fig. S8.**
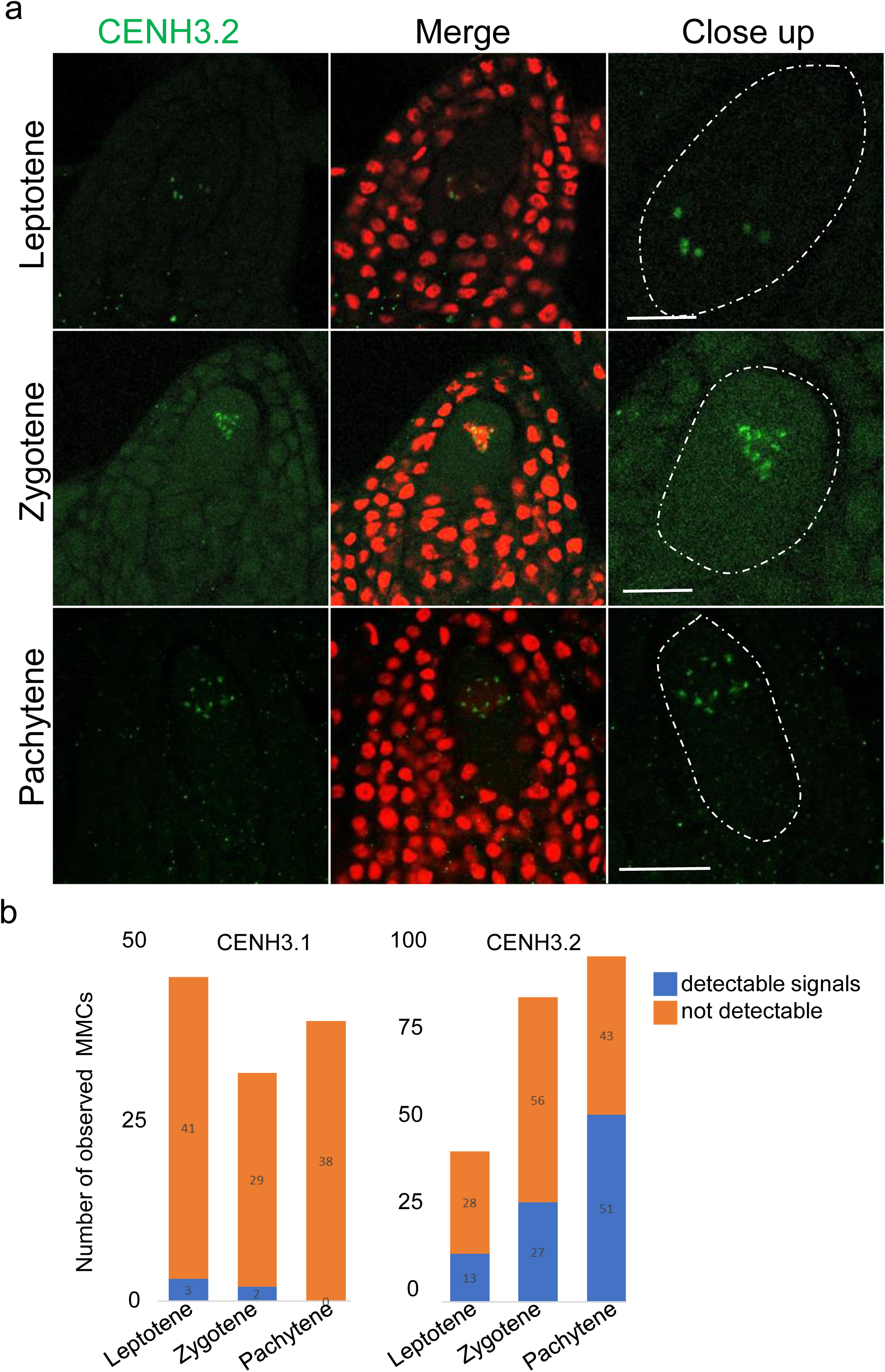
Distribution of CENH3.1 and CENH3.2 during female meiosis of cowpea tissue sections. CENH3.2 immunostaining signals (green) at leptotene, zygotene and pachytene (a). Nuclei were counterstained with propidium iodide (red). Frequency of observed CENH3.1 and CENH3.2 signals in leptotene, zygotene and pachytene cells (b). Bars equal 10 µm.

**Fig. S9.**
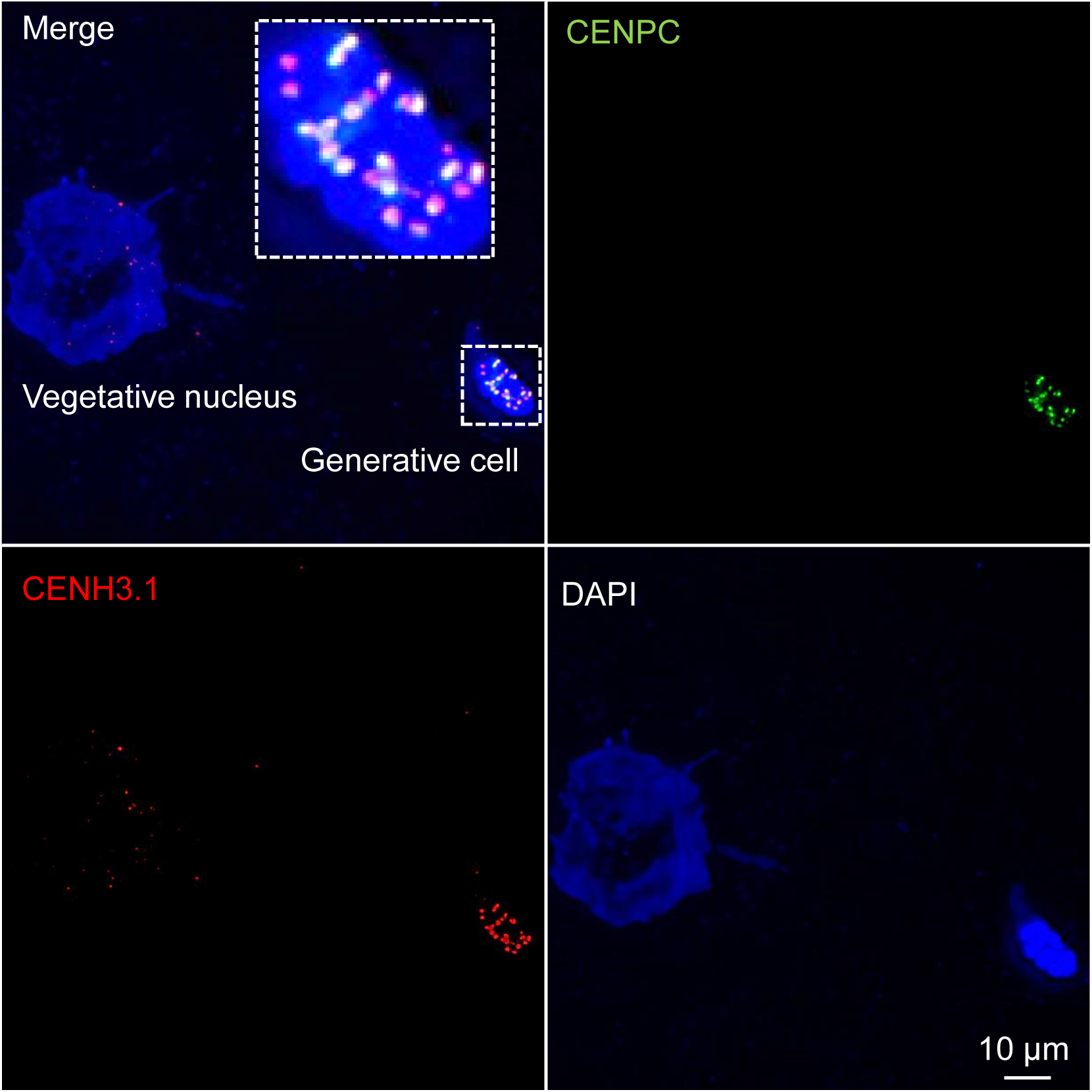
The generative nucleus of mature cowpea pollen shows colocalizing CENH3.1 (red) and CENPC (green)-specific immunosignals. The vegetative nucleus is free of centromeric immunomarks.

**Fig. S10.**
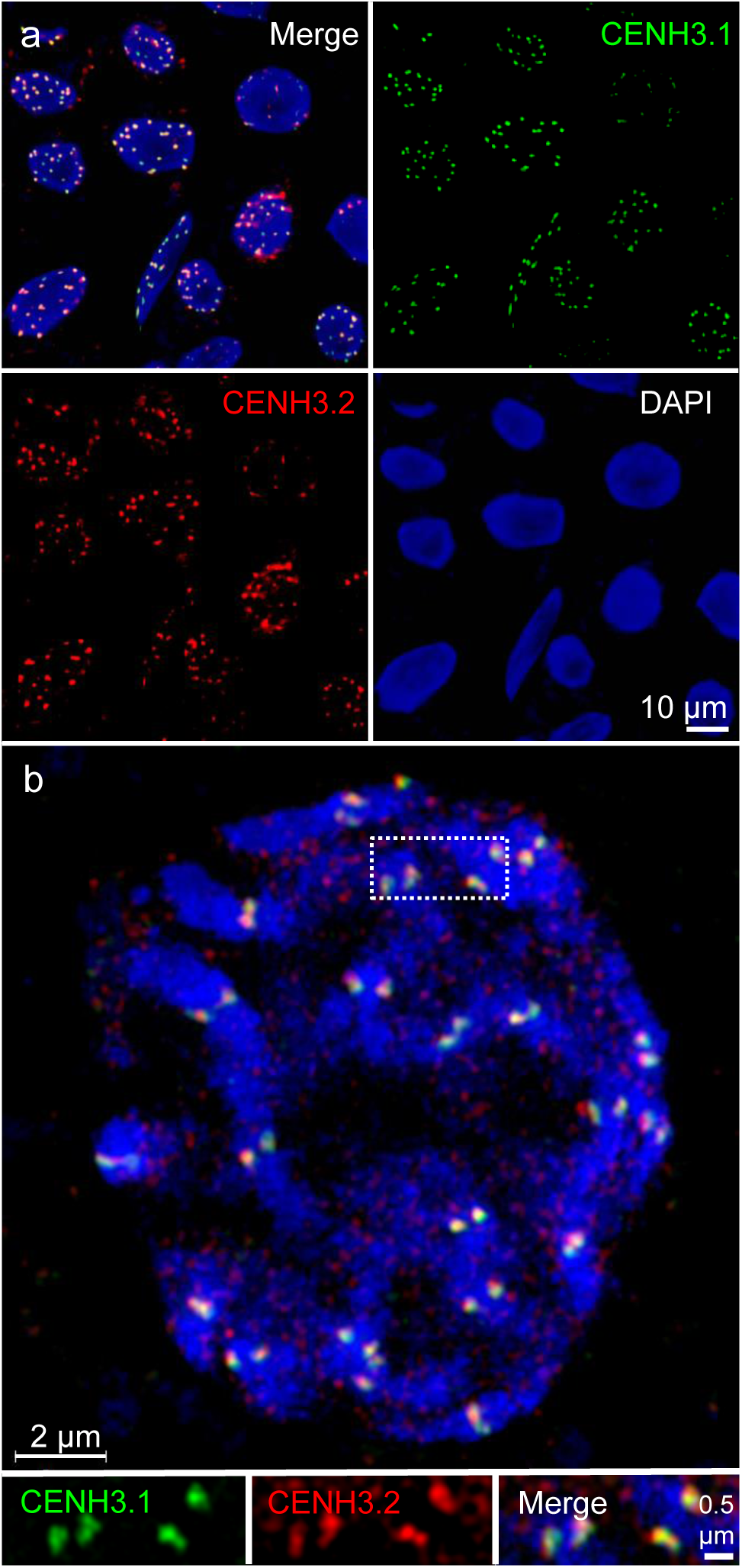
The centromeres of embryonic cells at early heart stage contain CENH3.1 and CENH3.2. CENH3.1 (green) and CENH3.2 (red) immunosignals colocalize in interphase (a) and prometaphase (b) cells. Structured Illumination Microscopy by SIM shows that both CENH3 variants occupy different centromeric subdomains (further enlarged inserts).

